# Twisted plywood-like tissue formation *in vitro*. Does curvature do the twist?

**DOI:** 10.1101/2023.09.04.556075

**Authors:** Barbara Schamberger, Sebastian Ehrig, Thomas Dechat, Silvia Spitzer, Cécile M. Bidan, Peter Fratzl, John W. C. Dunlop, Andreas Roschger

## Abstract

Little is known about the contribution of 3D surface geometry on the development of multi-layered tissues containing fibrous extracellular matrix components such as those found in bone. Here we elucidate the role of curvature in the formation of chiral, twisted plywood-like structures. Tissues consisting of murine pre-osteoblast cells (MC3T3-E1) were grown on 3D scaffolds with constant mean curvature and negative Gaussian curvature for up to 32 days. Using 3D fluorescence microscopy, the influence of surface curvature on actin stress-fiber alignment and chirality was investigated. To gain mechanistic insights, also MC3T3-E1 cells deficient in nuclear A-type lamins or treated with drugs targeting cytoskeleton proteins were used in our study. We find that wild type cells grow multilayered tissue with fibers predominantly aligned along directions of negative curvature, but where subsequent layers twist in orientation with respect to older tissues with time. Fiber orientation is conserved below the tissue surface thus creating a twisted plywood like material. We further show that this directional organization strongly depends on structural components of the cells (A-type lamins, actin and myosin). Our data indicate the importance of substrate curvature in the formation of 3D tissues and provides new insights into the emergence of chirality.

**Significance Statement:** Biological tissues (like compact bone) often consist of multiple fibrous layers which are staggered with a twisting angle relative to each other, thereby improving mechanical performance. The underlying principles of how such tissues are formed and what determines the fiber direction are still debated. Here we report the formation of a twisted plywood-like tissue grown *in vitro* on constant mean and negative Gaussian curvature substrates and present evidence that for tissue consisting of pre-osteoblast like cells, surface curvature is a main determinant for fiber orientation.

## Introduction

Structural biological materials, such as bone, wood, and cuticle, are composite materials (1, 2), and can consist of biopolymers such as cellulose, chitin and collagen. These polymers are long and thin, and form fibers or fibrils, which in turn pack parallelly into lamellae (1). Such lamellar structures are ubiquitous in Nature, as seen in compact bone (3), the exoskeleton of crustaceans (4), the cell wall of plants (5) and many more (1). These tissues typically consist of multiple layers of lamellae, giving rise to the so-called twisted plywood or helicoidal structure, in which the fiber direction changes from layer to layer. There are mechanical advantages for a natural material to have a twisted plywood structure. It results in isotropic material properties of a tissue formed from intrinsically anisotropic building blocks, but more importantly such twisted layers are excellent in resisting the propagation of cracks, which explains the high fracture toughness in so many biological materials (3, 6-9). A well-studied example is compact bone in which mineralized fibers of collagen type I build up ∼ 5 µm wide lamellae with an alternating twisting angle to form a plywood-like material with high toughness (9-13). Despite the well-known connection between structure and mechanical function, the processes causing the emergence of twisted plywood structures in Nature remain unclear. One theory suggests that self-assembly of these biopolymers into liquid crystalline phases gives rise to these structures (1, 14). Furthermore, it is known that cells can align themselves with each other over long distances as has been seen in studies of cells seeded on flat fibronectin patches of different shapes (15). Multilayered tissues are difficult to grow *in vitro* on flat surfaces in 2D, however for curved surfaces the situation changes, and thick tissues can form in which multiple layers of cells can be found. This is relevant of course to processes occurring on the highly curved surfaces found in trabecular (16, 17) and osteonal bone (18), where during bone formation a layer of osteoblasts forms new tissue. The presence of curvature, however, can in turn change the behavior of cells, modulating pattern formation and influencing the development of helicoidal tissues (19). To understand the development of such tissues it is thus necessary to determine the role of curvature on the long-range patterning of cells and the extracellular matrix that they produce.

Surface curvature has been shown to be an important geometric signal that plays a role in the fate and behavior of single cells and tissues (20, 21). For instance, mesenchymal stem cells (MSCs) seeded on convex semi-cylinders aligned preferentially along the zero curvature or axial direction, whereas MSCs seeded within concave semi-cylinders oriented towards the direction of highest curvature magnitude perpendicular to the cylinder axis (22). MSCs are also known to exhibit “curvotaxis” (23) or curvature-guided migration (22). On negative Gaussian curvature surfaces, they migrate along concave (valley-shaped) directions and on sinusoidal surfaces cells migrate towards the concave valleys (23, 24). Multi-cellular tissues formed by murine pre-osteoblast cells (MC3T3-E1) within prismatic pores show higher growth rates in concave compared to flat regions (25, 26). Tissues grown on triply periodic minimal surfaces (also with negative Gaussian curvature) display higher levels of osteogenic differentiation markers than tissues grown on control scaffolds, indicating that tissues also respond to surface curvature (27). An important hint towards the mechanism controlling this curvature sensing is the observation that a growing tissue has a shape that can be described by the Young-Laplace-Law (28). This link between pressure (mechanics) and curvature, suggests the importance of mechanical signaling in the sensing of curvature by cells and tissues, which is in turn further supported by computational modelling (29, 30). Although cellular mechanosensing clearly plays an important role in curvature driven cell migration and tissue growth (31, 32), the underlying biological mechanisms are not yet clear. Recent studies suggest the shape and the deformation of the nucleus (33, 34), the amount and arrangement of focal adhesion formation (23, 34) and the organization of the cytoskeleton (34) as potential key players in determining curvature sensing by cells.

An additional aspect that is required to understand how geometry influences the development of multilayered tissues is that cells may display preferential symmetries, preferring to orient with a left-or right—handed twist with respect to their surroundings. The actin cytoskeleton of single cells cultured on a circular patch exhibits a left-right asymmetry (35-37). Such chiral arrangements were also observed in cell monolayers confined to surface patches for several cell types (35, 38, 39). Remarkably, it has also been shown that interfering with certain actin associated proteins or tempering actin polymerization by drugs can reduce, eliminate or even reverse this effect in single-cells and cell-monolayers (35). Symmetry breaking in cell and tissue patterning can also be observed on hyperbolic surfaces, where pre-osteoblast-like cells were shown to self-organize into a left-handed chiral pattern (28). It is hard to pin down the ultimate cause of the asymmetric behavior of a growing tissue, as asymmetries occur across multiple length scales and may be causally related (40, 41). Single-cell experiments revealed that the asymmetry of the cytoskeletal proteins plays an important role in the polarity of cells, which in turn influences the asymmetry of tissues and organs (35-37).

Hence, it is possible that multilayered twisted plywood-like tissues develop through the complex interaction between mechanical signaling, curvature sensing and the innate asymmetry of the tissue producing cells themselves. To search for such interactions, investigation of tissues formed by cells needs to consider the 3D-nature of tissues and especially the effect of curvature on cells, as well as potential underlying cellular and sub-cellular mechanisms causing the emergence of chiral patterns on higher hierarchical levels. As tissue shape and thus also surface curvature changes during growth, curvature induced cell and tissue alignment may adapt with time. We explore the coupling between these aspects by growing multi-layered tissues produced by MC3T3-E1 pre-osteoblast cells on rotationally symmetric scaffolds of constant mean but negative Gaussian curvatures. Tissue orientation is quantified as a function of growth time by evaluating actin stress fiber and collagen fiber orientation with respect to the principal curvature directions.

Mechanistic insight is gained by comparing wild-type cells with cells exhibiting impaired nuclear mechanosensitivity (lamin A/C – deficient cells), as well as treatment for inhibited actin-polymerization (Latrunculin A), inhibited myosin II (Blebbistatin) and enhanced actin stress fiber formation (TGF-β1). By measuring how cell and tissue orientation evolves with time we start to unravel the complex mechanisms determining large scale tissue organization.

## Results

### A) Emergence of left-handed chirality in tissue grown on negative Gaussian curvature surfaces

To assess the role of surface curvature on the organization of tissues formed by pre-osteoblast cell cultures, we seeded MC3T3-E1 cells on two types of negative Gaussian curvature surfaces and investigated the development of tissue as a function of time. We used rotationally symmetric capillary bridges (28, 42) as one surface type and compare growth results on these surfaces with tissue growth within pores that are made by casting PDMS around a capillary bridge. In this way we can compare cell behavior on surfaces with identical Gaussian curvature, but with a mean curvature of opposite sign as well as with opposite signs of principal curvatures (see Material & Methods and Figure S14).

Before describing the orientation of tissues on these two types of surfaces it is useful to define clearly what we mean by tissue orientation and how it changes in subsequent tissue layers during growth. This is important as the growth directions of both surfaces are opposite, with outward growth on the capillary bridges and inward growth on the pore surfaces, making descriptions of chirality and its change with growth sometimes confusing. In a previous study (28) we observed that helical actin stress fibers form in tissues grown on capillary bridges and we define the helicity of these stress fibers in the same way, with left-handed or right-handed chirality of the helical structures observed. The difficulty is that when we make images of the growing tissues on the two surface types we do this from different directions with respect to the helical tissue (Figure 1 a, b). For example, on capillary bridges, a left-handed helical tissue, when observed from outside the bridge, would have an angle θ lying between 0° and 90° with respect to the negative x-axis of the image. For a tissue with the same helicity but growing within a pore, and observed from the inside, the angle θ would between 90° and 180° with respect to the negative x-axis of the image (Figure 1 a-d).

**Figure 1.**
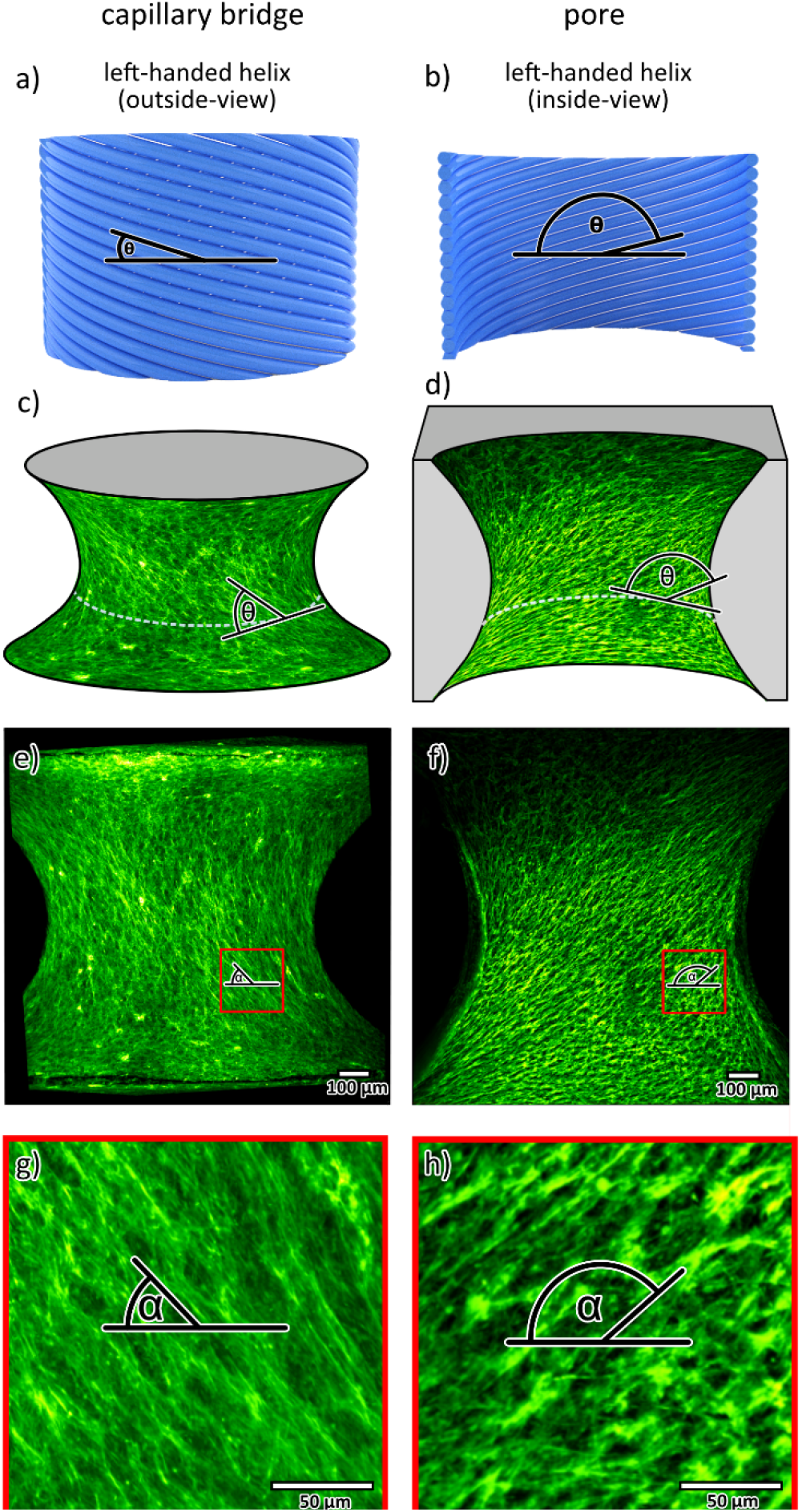
Definition of the fiber angle *θ* based on measurements using the outside-view (a) or the inside-view (b) of the same left-handed helix. MC3T3-E1 cells grown on a capillary bridge (c,e,g) and in an oppositely shaped pore (d,f,h) for 32 days and stained for actin (green). (a,d): 3D illustration of the grown tissue (PDMS scaffold shown in grey), the angle *θ* is measured in the 3D surface (0°: horizontal). (e,f): maximum intensity projections of the actin signal, indicating the fiber angle α. (g,h): zoom-in into the respective red indicated areas. Note that α denotes the fiber angle on the maximum projection image while θ is the fiber angle on the curved surface (see supplementary Figure S15). The two datasets are representative examples.

We first focus on the quantitative analysis of the actin fiber alignment on a capillary bridge (Figure 1 a,c,e,g) and a pore (Figure 1 b,d,f,h) after 32 days of culturing, which reveals the formation of helical stress-fibers with a left-handed chirality for both sample types. The capillary bridges have fiber angles *θ* smaller than 90° (Figure 1c) and the pores fiber angles *θ* higher than 90° (Figure 1d). The different angles measured for the two sample types are both indicating a left-handed chiral pattern. This is due to the differences in imaging conditions: The tissue grown on the capillary bridge is observed from the outside of the helix and the tissue grown in the pore is observed from the inside of the helix (see Figure 1a-d). This formation of a tissue with left-handed chirality after 32 days is consistent with previous reports on experiments performed on capillary bridges (28).

### B) Tissue twisting during growth

The observation described above raises several questions: (i) How consistent is the tendency towards the observed chirality? (ii) At which time point does the chiral pattern emerge and is the actin fiber alignment different in earlier culturing stages? (iii) What is the main factor determining final and potentially transient actin fiber orientations? To address these questions, we chose a time-series approach: Capillary bridges and pore samples were fixed at different time points of culturing (day 4, 7, 11, 16, 23, 32) and imaged with light sheet microscopy using a fluorescent actin stain.

For capillary bridges, we observed fiber angles *θ* distributed around 90° meaning stress fibers are oriented close to meridional directions (i.e. fibers are close to being parallel with the pore or bridge rotation axis). At day 4 the tissue orientation (*θ* -angle) is right-handed and mainly adopts angles higher than 90° 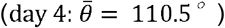 With ongoing culturing time 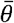 gradually decreases and adopts values below 90° after more than 16 days of culturing. This indicates the tissue gains a left-handed chirality during growth 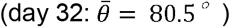, see Figure 2 c, e.

**Figure 2.**
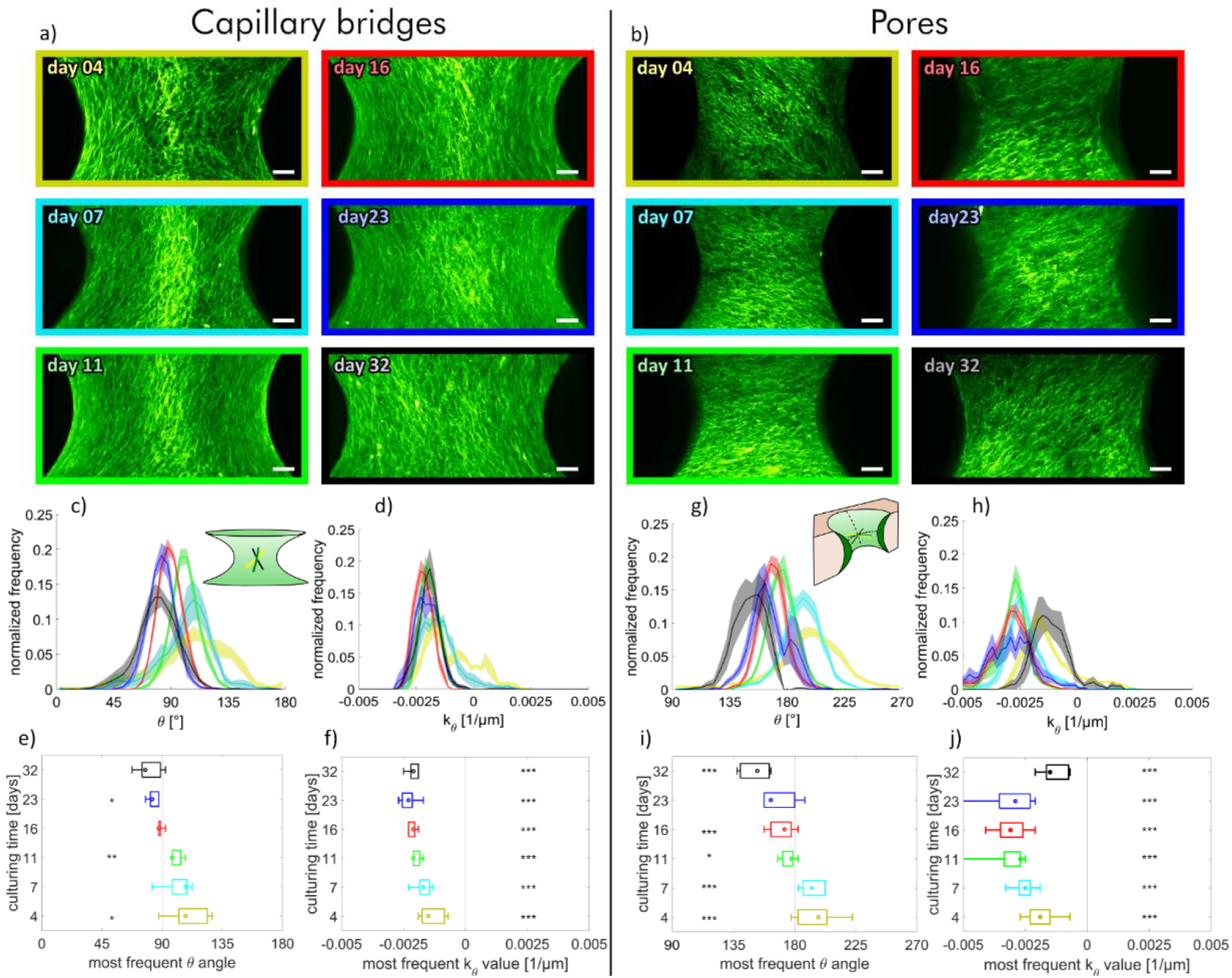
Evolution of the cell-orientation of tissues grown on (a) capillary bridges and (b) in scaffold-pores. Images are maximum intensity projections of samples stained for actin fixed at day 4 (yellow), 7 (turquois), 11 (green), 16 (red), 23 (blue) and 32 (black), respectively (scale bars: 100 µm). (c, g): histograms of the actin fiber angle distribution averaged over all samples with the same culturing time; (d,h): histograms of the measured normal-curvature along the *θ*-direction (*k*_θ_) averaged over all samples with the same culturing time. Shaded regions refer to the standard error of the mean for each bin; (e,f,i,j) median, interquartile intervals and range of the peak *θ*-angles and *k*_*θ*_ values are as a box and whisker plot. Single sample histograms of all samples are shown in the SI Figure S1 and S2. The number of independent samples for every time point are: n_CB day 4_ = 5, n_CB day 7_ = 5, n_CB day 11_ = 5, n_CB day 16_ = 5, n_CB day 23_ = 5, n_CB day 32_ = 5, (g-j) n_Pore day 4_ = 29, n_Pore day 7_ = 17, n_Pore day 11_ = 17, n_Pore day 16_ = 16, n_Pore day 23_ = 7, n_Pore day 32_ = 7. Statistical significance between most frequent θ-angles and 90° (e) or 180° (i) or most frequent curvature *k*_θ_ and *k*_θ_*=*0 1/µm (f, j) is indicated with p < 0.05 (*), p < 0.005 (**) and p < 0.001 (***).

In contrast to tissue growing on capillary bridges, tissue grown in pores align consistently around the equatorial direction with angles being centered around *θ* = 180° (or 0°). However, in a similar manner to capillary bridge tissues, at early stages (e.g. day 4) the tissue has a right-handed chirality 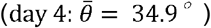 and also gradually shifts towards a left-handed chirality with increasing culturing time 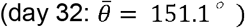 (see Figure 2 g and i).

Negative Gaussian curvature implies that any line on a surface, and thus the cells and stress-fibers, can have positive, negative, or even zero curvatures depending on their local orientations. As the sign of curvature is somewhat arbitrary, we define a concave direction as one which has a negative normal curvature in that direction. An example would be the meridional direction on an inward waisting capillary bridge. A convex direction would have positive normal curvature such as the equatorial line around the waist of a capillary bridge. Clearly on the pores the signs of principle curvature swap with respect to the capillary bridges, meaning the equatorial lines are concave and meridians are convex using this definition.

We used 3D models of the experimental surfaces to evaluate the normal curvature *kθ* of the fibers. *k*_*θ*_ was found to be relatively constant for the capillary bridges over the entire culturing time with typical values of -0.002 µm^-1^ (Figure 2 d and f), indicating the tissue’s tendency to consistently align along slightly negative curvature (concave) directions. Surprisingly in tissues grown in pores, an alignment along concave directions was also observed (Figure 2 h and j). The curvature distributions of fibers within the pores is broader than those of the capillary bridges. An interesting similarity is that for both sample types the curvature at day 4 is similar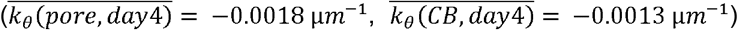] On both surface types, the normal curvature *k*_*θ*_ becomes more and more negative during growth, with only the last data points of the tissues grown in pores (day 32) showing a slight increase in curvature. For all samples and time points, *k*_*θ*_ was highly significantly different from 0.

The *θ*-values reflect the actin alignment on the tissue surface at different time points allowing us to hypothesize about changes in tissue orientation with growth. In both sample types the actin fiber orientation changes from a right-handed helix to a left-handed one during growth. This implies a negative twist for both tissues grown in pores and on capillary bridges. This consistent twisting of the helical orientation of the actin stress-fibers as a function of growth time potentially suggests that multiple cell layers form a twisted plywood structure. An alternative explanation is that cells in the bulk re-orient themselves with time and hence no correlation between fiber orientation on the surface and in the bulk should be found. To test this, we performed an analysis of the fiber orientation as a function of depth profiles of the tissues.

### C) Conserved fiber orientation of the tissue

As the results in Figure 2 indicate that orientation of newly deposited tissue changes with growth, we next determined how the tissue is organized below the surface. We define any change in helicity with growth as the twist, with a positive twist meaning the angle θ increases as the tissue grows (outwards (capillary bridge) or inwards (pore)) and a negative twist means the angle θ decreases with increasing layer thickness (Figure 3, black arrows). To do this we measured the actin fiber orientation at the neck region of each image of the image stacks derived from the light sheet data sets after 16 days of culturing (see Figure 3). This time point was chosen as this is the first time point where a flip towards a left handed-chirality is shown (see Figure 2). On the capillary bridges the tissue below the surface (∼60 µm) aligns in a right-handed chiral pattern with fiber angles above 90° whereas the surface tissue exhibits angles below 90° (Figure 3e). This is consistent with the negative twist observed in surface actin orientation from the time series experiments described above. A similar twisted plywood like organization is also observed in the pore samples (Figure 3f). The two datasets shown in Figure 3 are representative examples. For both sample types, nearly all of the obtained datasets showed a comparable behavior.

**Figure 3.**
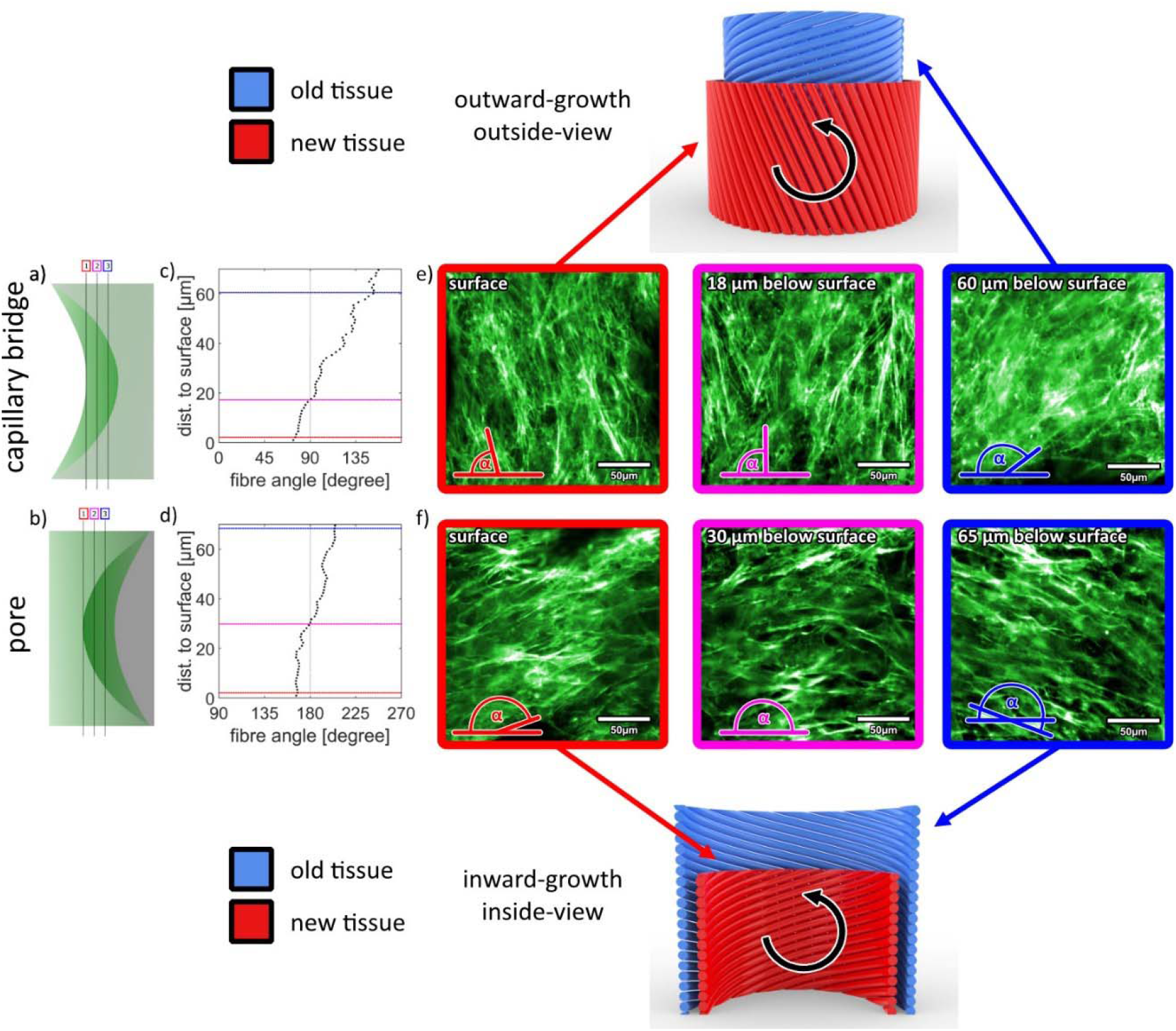
Actin fiber orientations at different depths below the tissue surface of MC3T3-E1 cells grown for 16 days on a capillary bridge (a,c) or inside a pore scaffold (b,d). Each data point represents the result of a fiber-orientation measurement of a single slice within a region of 200 x 200 µm^2^. (e,f) Fluorescence images showing the actin fiber arrangement in selected slices (frame color corresponds to line in plot, brightness/contrast is adjusted to compensate for signal loss below the surface). The sketches illustrate the spatial and temporal development of the fiber orientations for both setups. The black arrows in counter-clockwise direction indicate that in both cases a negative twist is present with ongoing tissue growth. The two datasets are representative examples; results for other samples can be found in the SI Figures S3, S4 and S12. A video of a depth profile is shown in the supplementary files.

### D) Collagen fibers follow actin stress fiber orientation

As MC3T3-E1 cells treated with ascorbic acid (included in our growth medium) are known to synthesize highly-ordered collagen fibrils (43), we next investigated the collagen fiber alignment inside the tissue. Confocal and second harmonic generation depth scans were performed to obtain the orientation of collagen and actin fibers of tissue grown on capillary bridges for 16 days. This analysis led to two main results: (i) In the outer layers of the tissue actin is present but no collagen (Figure 4 c, red box) while deeper regions contain both actin and collagen (Figure 4 c magenta and blue). The offset of the collagen signal relative to the actin is 5-10 µm (Figure 4 a). (ii) From Figure 4 b as well as from the selected slices shown in 4 c, it is clearly visible that actin and collagen fibers are oriented in similar directions. This is observed throughout the entire sample thus indicating a consistent co-alignment of cellular and extracellular components (see also supplementary Figures S12 and S13 and the corresponding videos in the supplementary files).

**Figure 4.**
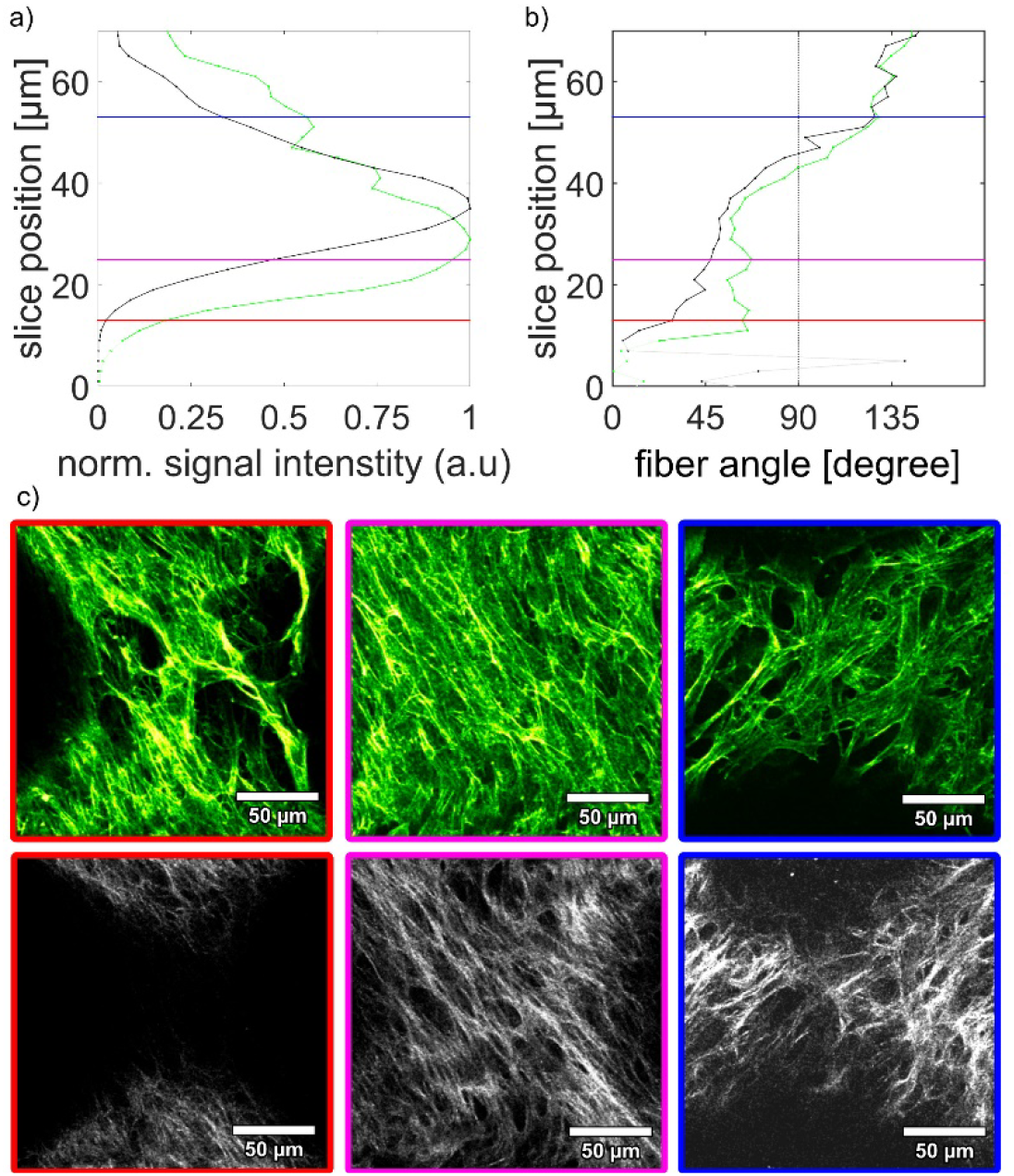
Co-alignment of actin and collagen fibers. a) intensity profiles of the actin (green) and collagen (black) image stacks (moving from outside the sample (0) 70 µm deep into the sample); b) results of the fiber angle evaluation for actin (green) and collagen (black) – very low intensity regions are shown in light colors; vertical lines correspond to the selected image slices shown in c): actin (upper panel) and collagen (second harmonic generation signal, lower panel). Image brightness and contrast settings were adjusted for visibility. More datasets can be found in the SI Figure S5 and S13.

### E) Investigating the role of nuclear mechanosensing, cytoskeleton and contractility

To obtain mechanistic insight into the change of the cellular orientation during culture, we first tested the impact of lamin A/C deficiency associated with impaired mechanosensing of the nucleus (44). Therefore, lamin A/C-deficient MC3T3-E1 cells (LAC-ko) were generated using the CRISPR/Cas9-system (for details see supplemental information and SI Figure S6) and were grown on capillary bridge samples. At the beginning of cell culture (day 7), LAC-ko cells show a right-handed chirality with *θ* – angles clearly larger than observed in wild type (WT) cells. In contrast to the WT-cells, there was no twist of the growing tissue towards a left-handed chirality (see Figure 5). Instead, the tissue remained the original right-handed chiral pattern even after 32 days of tissue culture. The development of *k*_*θ*_ was remarkably similar in both groups (Figure 5 d). Visually, the tissue formed by the LAC-ko cells appears to be much looser compared to the WT tissue (Figure 5 e, f).

**Figure 5:**
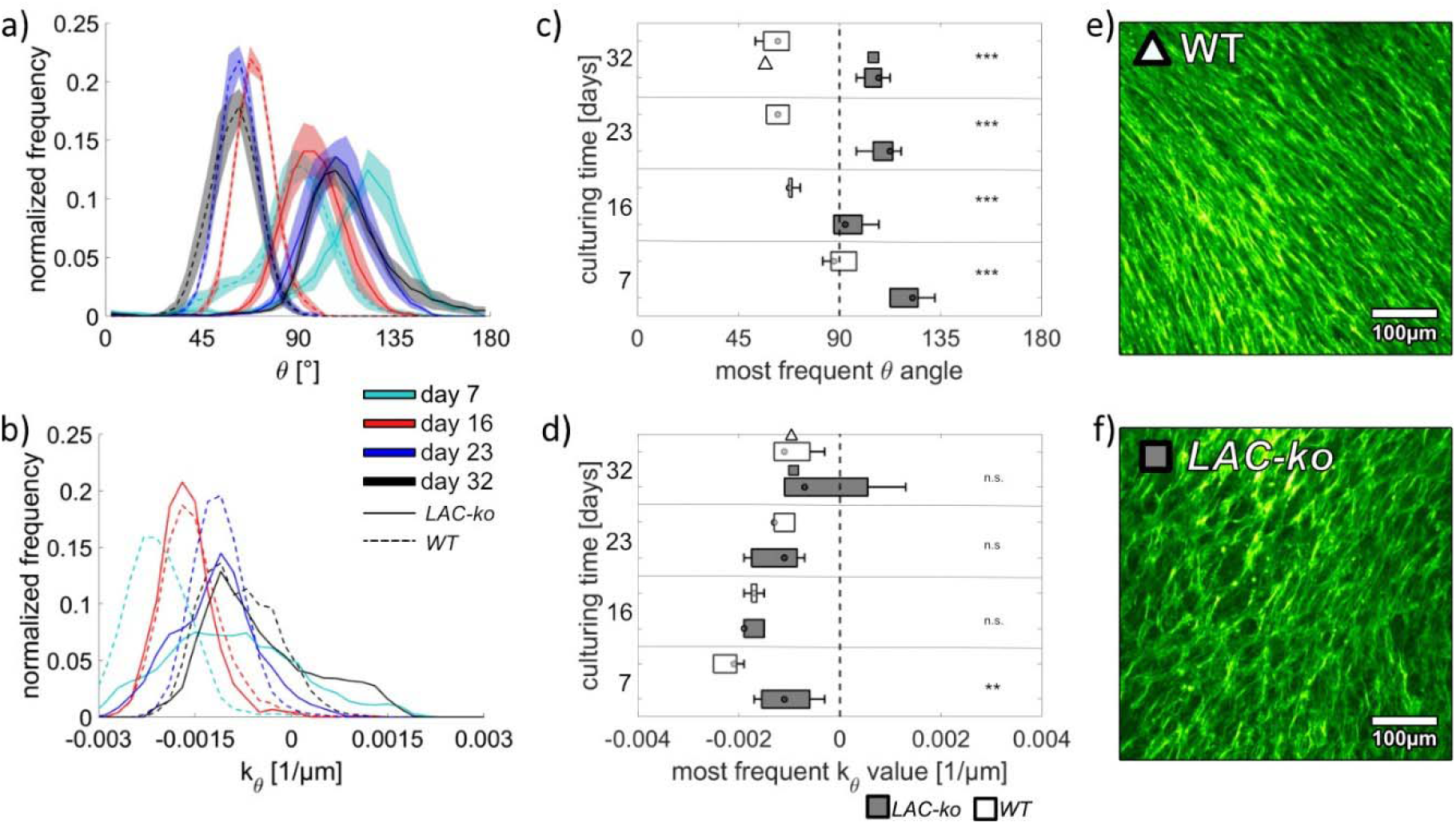
Cell-orientations of lamin A/C deficient MC3T3-E1 cells (LAC-ko, solid lines, grey boxes) compared to wild type (WT, dashed lines, empty boxes) grown on capillary bridges. (a): histograms of the actin fiber angle distribution averaged over all samples with the same culturing time. Shaded regions refer to the standard error of the mean for each bin; (b): histograms of the measured normal-curvatures along the fiber direction averaged over all samples with the same culturing time. (c) Mean and standard distribution of the peak theta-angles and (d) the curvature along the fiber direction. (e,f): Two examples of the center-regions after 32 days of culturing. Symbols are used to indicate the respective values in (c) and (d). Sample numbers: n=5 for each group. Histograms for the single samples can be found in SI Fig. S7. Statistical significance between WT and LAC-ko for each time point is indicated with not significance (n.s.), p < 0.005 (**) and p < 0.001 (***).

To investigate the effect of the cytoskeleton and contractility, MC3T3-E1 cells grown on capillary bridges were treated with either Blebbistatin (inhibition of myosin II), TGF-β1 (stabilization of actin stress fibers; decreased motility) or Latrunculin A (LatA; inhibition of actin fiber assembly). Samples were fixed on day 7 and day 32. Blebbistatin treated samples show at day 7 and day 32 a left-chiral alignment while the controls exhibit a flip from right to left during this period (Figure S8). In contrast, TGF-β1 treatment leads to right chiral arrangement at both time points indicating a conservation of the right-handedness (Figure S9). With LatA treatment a non-significant trend towards smaller θ-angles was observed for day 7 while θ was significantly smaller compared to control at day 32, where also a left-handed chirality was observed as in the controls (Figure S10). To exclude the effect of DMSO, which is used as a solvent for Blebbistatin and LatA, additionally samples were treated with DMSO only. In this case no significant effect on the tissue organization compared to the control samples was observed (Figure S11).

## Discussion

There are three main aspects that we learn from observing the time and depth dependent organization of tissues grown on negative Gaussian curvature surfaces:

i. We observed that during cell culture there was a consistent change in the orientation of the surface tissue layer relative to the layer directly beneath it. This suggests that cells sense underlying tissue orientation and then produce a new orientation with a negative twist. Depth profile measurements demonstrated that this arrangement is preserved in the bulk, thus resulting in tissue organized similar to parts of a “twisted plywood like tissue” (1). Our results further suggest that the cells located below the surface layer (∼10 µm below the tissue surface) lay down a collagen matrix which is co-aligned with the cell’s actin cytoskeleton, thus hinting at the mechanism giving rise to extracellular matrix organization in such tissues.
ii. The normal curvature along the main actin fiber direction was always negative (concave). Despite the change in mean surface curvature during growth, the normal curvature remained close to - 0.002 µm^-1^ for both capillary bridges and pores (with a larger scatter in the data for the latter). This suggests a mechanism by which a twist in orientation appears in order for the growing tissue layer to retain this preferred curvature. A consequence of this is that fiber orientation in a growing tissue is strongly mediated by changes in surface geometry due to growth.
iii. The results reveal a consistent change in tissue chirality. While in early tissue culture stages actin stress fibers and collagen are oriented in a right-handed chiral structure (capillary bridges: 90 < *θ* < 180, pores: 180 < *θ* < 270), after around two weeks of tissue culture a left-handed chirality is adopted. This behavior revealed to be remarkably similar for both sample types. It is still unclear how the first layer of tissue becomes ordered in the early stages of growth and what triggers the observed changes in orientation.

We hypothesized that the underlying mechanism of tissue patterning is linked to the mechosensitivity of cells. Compared to the wild-type tissues, tissues formed by a lamin A/C-deficient cell-line as well as those of tissues treated with drugs to modulate actin-myosin contractility displayed modified twisting behavior. After short culturing times, Lamin A/C-deficient cells and cells treated with Blebbistatin or TGF-β1 form spiral patterns, but different to wild type cells, longer culturing times do not lead to a change in chirality and thus to the absence of tissue twisting. This might indicate that especially the extent of twisting is sensitive to disturbances in the actin skeleton and nuclear lamins. These findings agree with recent studies showing a connection between curvature, matrix stiffness, actin-cytoskeleton and lamin A/C levels. For example, lamin A/C levels increase with curvature as well as with surface stiffness, the latter being dependent on actomyosin (45-48). Although the exact mechanism underlying cell layer alignment and twisting are not yet known, it is possible that Yes-associated protein 1 (YAP)-signaling is involved, which functions in mechanotransduction and bone development and is decreased by lamin A/C deficiency (45, 49, 50). Additionally, impaired cell migration, which has been observed in cells lacking lamin A/C (51), might also be accountable for the defective twisting behavior in the LAC-ko cells. Cell migration plays an essential role in the ordered formation of bone tissue (52) and is influenced by surface topology (21, 53, 54).

Low doses of LatA were shown in literature to result in inverted chirality of fibroblasts in single cell and tissue sheets (35). This was not observed in our study on MC3T3-E1 cells. The drug however led to a more pronounced clockwise rotation of the tissue which is observed by smaller θ-angles at the end of the culture compared to the control samples. The inhibiting effect of TGF-β1 on the twist might be explained by a higher contractility of the cells as well as their decreased motility (55). Surprisingly, the decrease of contractility by Blebbistatin leads already at the beginning of the tissue formation to a left-handed orientation, which continued to be so until the end of the culture. Although the specific mechanisms remain unknown, these data suggest that the emergence of chirality and tissue twisting is under cellular control and is influenced by nuclear mechanosensing and actin skeleton properties.

As the chosen MC3T3-E1 cells are preosteoblastic, it is remarkable that the observed tissue twisting presents similarities of what is found in bone. Osteons located in human long bone for example exhibit a typical lamellar plywood-like structure with around 5 µm thick lamellae of mineralized collagen fibers (9-13). Despite the different tissue compositions (a highly mineralized collagen extracellular matrix vs. a loosely packed cell-collagen tissue) it is likely that our *in vitro* results showing the emergence of a chiral tissue structure are triggered by similar processes responsible for the twisted plywood structure in the *in vivo* case.

We previously demonstrated that tissue grown on capillary bridges adopts shapes known from fluids with isotropic surface tension, thus forming constant-mean curvature (Delaunay) surfaces (28). Further theoretical work investigated how the presence of fibrous tensile elements that follow geodesic lines on the surface, may modulate tissue shape, as it might be the case for highly aligned and orientated cell systems (56). For the fiber-angles observed in experiments, macroscopic surface geometries were remarkably similar to the Delaunay surfaces. The main difference is that fiber angles can strongly modulate internal tissue pressure, thus strengthening the role of mechanosensing in tissue orientation.

Neville cites three possible mechanisms leading to the formation of twisted plywood tissues in biological materials (1): Self-assembly of molecules, directed assembly by cellular mechanisms or mechanical reorientation. Our results suggest that the development of a twisted plywood like tissue relies on a mixture of the first and second mechanisms, in which the cells self-organize into ordered arrays resembling liquid crystals, and then control collagen alignment. Although Neville’
ss work considered molecules which passively self-assemble, also contractile forces mediated by the cell’s cytoskeleton may be considered in growing biological tissues (29). Further data suggests that cells may also spontaneously align like liquid crystals on a surface (15, 57, 58). Our data enables us to hypothesize a mechanism for tissue organization. We however cannot say why the first layer of cells has the orientation it does as in principle both left- and right-handed chirality of tissue layers should allow cells to experience identical curvatures. Initial asymmetry is likely to arise from the well-known intrinsic cell-chirality (35, 59). Once the first layer of cells has formed, subsequent cell layers are deposited in manner such that they can keep aligned with the preferential curvature direction giving rise to a twist relative to the layer below. As the tissue thickens collagen is produced in an already helical environment of cells and tissues, thus freezing this helical arrangement into place. This is in contrast with the hypothesis that procollagen molecules form a liquid crystal phase and thus shape the emerging tissue in bone (60). This alternative hypothesis is based on the observation that also in the absence of cells under well controlled conditions collagen can spontaneously form cholesteric liquid crystalline phases similar to a twisted plywood arrangement. Since it remains open to which extend this process may take place in a more complex environment, our results do not dispute the importance of self-organization processes of collagen but show that also processes at the level of cells can give rise to the emergence of twisted helices. It is becoming clearer that cellular control plays a crucial role, however we still do not understand how cells sense curvature although data give some hints as to the importance of mechanosensing. The hypothesis that the twist in tissue orientation is linked to changing curvature opens up some new predictions, which potentially can be tested experimentally. During remodeling of cortical bone, osteoblasts follow the cutting cone of osteoclasts depositing oriented tissue in the osteoclasts wake. If surface curvature indeed mediates cell and collagen orientation, then the speed of cell migration, and osteoclast resorption would also play an important role in tissue orientation, perhaps explaining the variation in osteonal structures seen *in vivo* (61).

In conclusion, the presented results reveal the importance of surface curvature for tissue alignment. Curvature sensing, mediated by actin-myosin cell-contractility and the nuclear lamins enables cells to align in particular curvature directions which change as growth proceeds. This change results in the formation of a layer of twisted plywood-like tissue. We demonstrate that such tissues can be formed *in vitro*, this alone gives us a new model system to help understand how complex multi-functional materials are formed by fibrous building blocks and to potentially control and optimize tissue orientation in the future.

## Materials and Methods

### Manufacturing of PDMS capillary bridges

Capillary bridges were fabricated according to the routines described previously (28, 42): For the fabrication of the scaffolds, the Sylgard 184 Elastomer Kit (Dow Corning, USA) was used with a mixing ratio of ten parts base with one part curing agent (weight : weight). After mixing the base with the curing agent, PDMS was degassed in vacuum for 20 min. Using a syringe pump (New Era pump systems, NY, USA), the PDMS was transferred to cylindrical aluminum pillars facing to each other with a radius of 1 mm. The amount of liquid dispensed on the pillars corresponds to the volume of the final capillary bridges and therefore was adjusted for the individual capillary bridge sizes. The pillars were moved together so that the PDMS liquid droplets got in contact and a liquid bridge formed between two pillars. The distance between the pillars was adjusted to 1.25 mm, which corresponds to the height of the resulting capillary bridges. The capillary bridges were cured at 120°C for 20 min.

### Manufacturing PDMS pores with negative Gaussian curvature

The capillary bridges manufactured above, were furthermore used as templates for pores with a negative Gaussian curvature. A thin layer of PDMS (see mixing procedure in the last paragraph) was applied on a four inch silicon wafer by spin coating (1100 rpm, 1 min, spin coater, Laurell Technologies Corporation, PA, USA). The capillary bridges were placed with a tweezer on the wafer and cured for 2 h at 80°C. For the casting of PDMS a method described in (62) was used: The wafer with the capillary bridges was treated with oxygen plasma (50 W, 1 min, Emitech K1050X) and afterwards directly placed in a bath of 100% ethanol for 30 min in vacuum and dried for 30 min at 80°C. A PMMA frame was glued on the wafer surrounding the capillary bridges and filled with PDMS while avoiding complete coverage of the capillary brides. Finally, the samples were placed in vacuum for 20 min and cured at 80 °C for 2 h. The PDMS with the pores was manually detached from the capillary bridges and cut into an appropriate size for cell culture. A graphical description can be found in the supplementary information S14.

### Functionalization of PDMS scaffolds with PDMS

The surfaces of the PDMS scaffolds were functionalized using a combination of plasma treatment, (3 aminopropyl) triethoxysilane (Sigma-Aldrich Chemie GmbH, Steinheim, Germany) and glutaraldehyde (Carl Roth GmbH + Co, Karlsruhe, Germany) to covalently bind fibronectin (Sigma-Aldrich Chemie GmbH, Steinheim, Germany) to the surface. The method was adapted from (63) and described in detail in (28).

### Cell line

For all experiments, except the lamin A/C deficient and the collagen imaging, MC3T3-E1 cells, were used. For the lamin A/C deficient experiments, MC3T3-E1 LAC-ko and MC3T3-E1 cells were provided by the Ludwig Boltzmann Institute of Osteology. This MC3T3-E1 cells were also used for imaging the collagen organization. LAC-ko cells were generated using the CRISPR/Cas9-system (for details see supplemental information and SI Figure S6).

All cells were tested regularly for mycoplasma contamination and the MC3T3-E1 cells were authenticated by STR analysis (Microsynth, Balgach, Switzerland).

### Cell culture

Murine pre-osteoblast cells MC3T3-E1 were seeded on the scaffolds with a density of 10^5^ cells/cm² and cultured using α-minimum essential media (α-MEM) (Sigma-Aldrich Chemie GmbH, Steinheim, Germany) supplemented with 4500 mg/L D-(+)-glucose (Sigma-Aldrich Chemie GmbH, Steinheim, Germany), 10% fetal bovine serum (FBS) (gibco, Life Technologies Limited, Paisley, UK), 50 µg/mL L-ascorbic acid (Sigma-Aldrich Chemie GmbH, Steinheim, Germany) and 1% penicillin-streptomycin (Sigma-Aldrich Chemie GmbH, Steinheim, Germany). The samples were incubated at 37°C and 5% CO_2_ in humidified atmosphere. The scaffolds were transferred into new plates every 7 days starting from day four. The cell culture media was exchanged every two to three days beginning at day four unless otherwise stated.

### Blebbistatin / TGF-b1 / Lat A

To change the contractility of the cytoskeleton the treatment of the samples with three different drugs was performed starting at day four after seeding until the end of the experiment: myosin-II inhibitor Blebbistatin to inhibit the myosin-actin contractility; TGF-β1 to enhance the contractility and stabilize the cytoskeleton or Latrunculin A to inhibit actin assembly. (-)-Blebbistatin (Sigma-Aldrich Chemie GmbH, Steinheim, Germany) was added to the growth media at a final concentration of 2 µM with 0.1% dimethyl sulfoxide (DMSO) (Sigma-Aldrich Chemie GmbH, Steinheim, Germany). The recombinant human TGF-β1 (Invitrogen, MD, USA) was added at a final concentration of 1 ng/mL. The actin monomer binding toxin Latrunculin A (Millipore, Darmstadt, Germany) was added at a final concentration of 20 nM with 0.1% DMSO. To exclude the effect of the solvent DMSO, additional samples were treated with 0.1% DMSO in the same period.

### Fluorescent staining and imaging

The tissue was fixed at the end of each experiment with 4% paraformaldehyde in PBS (Alfa Aesar, ThermoFisher GmbH, Kandel, Germany) for 5 min at room temperature followed by thoroughly washing using 1x Dulbecco’s Phosphate Buffered Saline (DPBS) (Sigma-Aldrich Chemie GmbH, Steinheim, Germany). The tissue was permeabilized with 1% Triton X-100 (Sigma-Aldrich Chemie GmbH, Steinheim, Germany) between three hours to overnight at 4°C and washed extensively with 1x DPBS. To visualize F-actin, samples were stained for 90 min with 1.65 x 10^−7^ M Alexa Fluor 488 phalloidin (Invitrogen, Life Technologies Corporation, Oregon, USA) in 1x DPBS. The tissue was again washed with 1x DPBS and to visualize the cell nuclei, the tissue was incubated for 5 min with 1 µM TO-PRO-3 iodide (Invitrogen, Life Technologies Corporation, Oregon, USA) followed by washing with 1x DPBS (nuclei data not shown). Prior to the imaging of the pore samples, after fixation and staining, the samples were cryo-embedded in OCT embedding matrix (Cellpath, Newtown, UK) and opened by cuting into half, followed by embedding 1% low melting agarose (Carl Roth GmbH + Co, Karlsruhe, Germany) for imaging at room temperature. For fluorescence imaging of capillary bridges and opened pores a Zeiss Z1 light sheet fluorescence microscope was used. The fixed tissue was placed in the imaging chamber filled with deionised H_2_O. A Plan Apochromat 20x/1.0 Corr DIC water immersion objective lens was used and a 488 nm and a 633 nm excitation laser. To image F-actin, the SBS LP560 beam splitter was employed. 3D data were generated by performing z-stacks with 0.63 µm lateral pixel size and 1.2 µm z-step size. For further analysis maximum intensity projections (MIP) of the stacks were used. Due to the attenuation of the fluorescence signal from regions inside the tissue, actin fibers on the surface are distinctly brighter than those below. Hence in the MPIs mainly the tissue surface is visible. For imaging collagen with two-photon excitation, a SP8 (Leica Microsystems GmbH, Wetzlar, Germany) confocal laser-scanning microscope equipped with pulsed tunable laser (Maitai, Newport Corporation, California) was used. The excitation wavelength was set to 910 nm while acquiring the emission signal in a spectral window between 430 and 470 nm. The Alexa Fluor 488 phalloidin signal was also acquired for each slice. Image stacks were obtained with a voxel size of 0.606 x 0.606 x 2 µm. The images were acquired using a Fluostar VISIR 25x/0.95 WATER objective.

### Analysis of fiber alignment

The actin fiber alignment was derived from light sheet microscopy fluorescence data of fixed and stained samples. Due to the anisotropic resolution, instead of using full 3D data, our results are based on maximum intensity projections (MIPs) as described above. In short: First, MIPs of the fluorescence images were then performed and the fiber direction in the projection plane was derived (α). To calculate the fiber angle on the sample surface (*θ*), vectors representing the fibre orientation were projected on rotational symmetric representations of the scaffold using in-house developed Matlab routines (Mathworks Inc., MA, USA, v 2020a). Edge regions as well as regions more than 300 µm above/below the neck center were excluded from the evaluation to avoid boundary artefacts. The routines for the fiber angle alignment are based on the work published in reference (28). In ∼80% of the capillary bridge samples and ∼20% of the pore samples it was possible to also obtain *θ*angles from the backside or second sample half respectively. In this case data from both measurements are included in the respective histograms. For every location where *θ* was evaluated, also *k*_*θ*_, the normal curvature in *θ*.-direction was derived. This was done by first calculating the principal curvatures (largest and smallest curvature) at the location and then using the Euler theorem of differential geometry to derive the curvature in the direction of *θ*.

A more detailed description of the evaluation procedure can be found in the supplementary information and Figure S15.

### Statistical Analysis

Statistical analyses were performed using Matlab. To compare two sample cohorts, the most frequent *θ* or *k*_*θ*_ value of every sample (histogram peak position) served as input data for a two sample t-test. When testing whether *θ* or *k*_*θ*_ are significantly different to a given value, a one sample t-test was chosen. Differences of p<0.05 are considered as significant.

## Supporting information

zStack_actin_collagen

wildtype_actin

wildtype_collagen

## Acknowledgments

The authors thank Peter Steinbacher for his support in sample preparation. B.S. thanks Daniel Hoeckner for his support in image analysis.

