## Supplementary Information for "Twisted plywood-like tissue formation *in vitro*. Does curvature do the twist?"

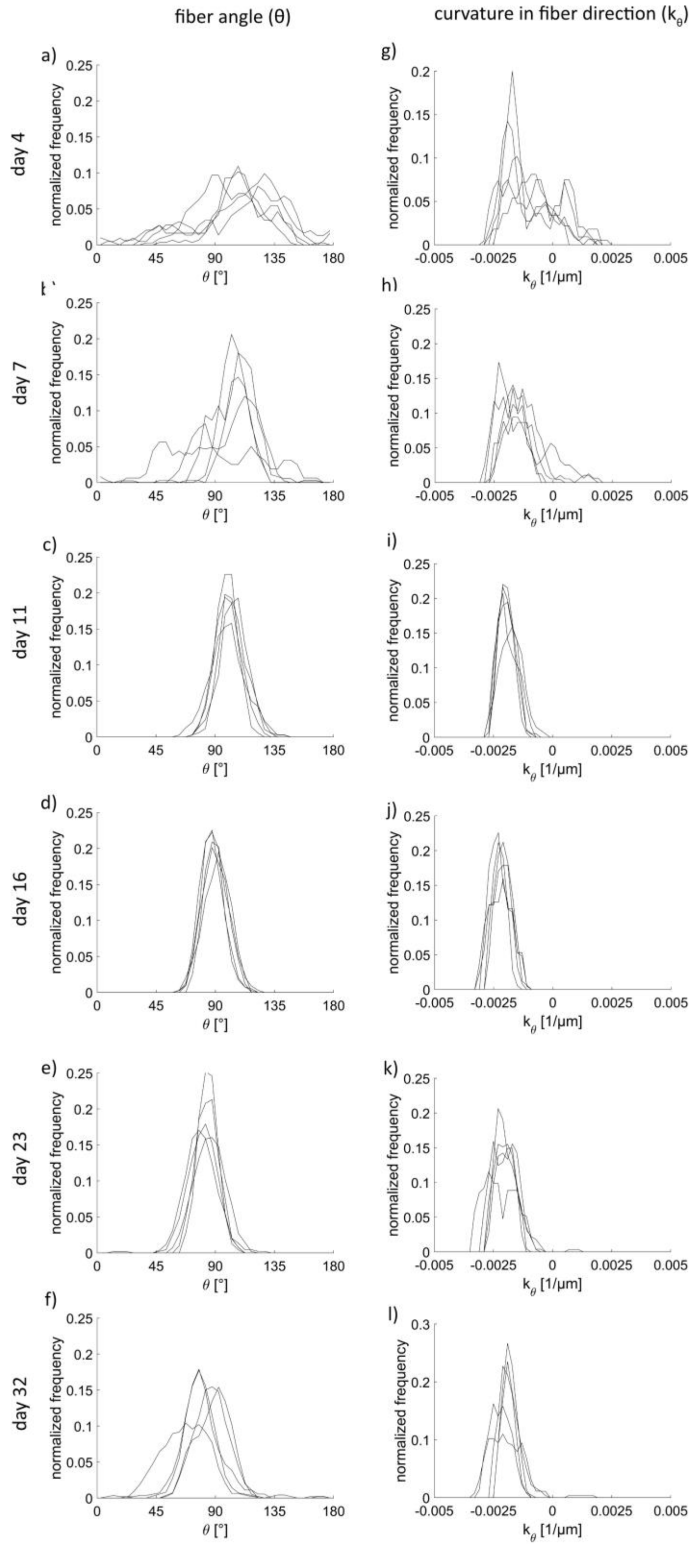


S1: add Fig 2: Angle distribution (a-f) for actin and distribution for curvature along theta (g-l) on capillary bridges at different timepoints. Each line indicates one sample. Each timepoint consists of five repetitions.

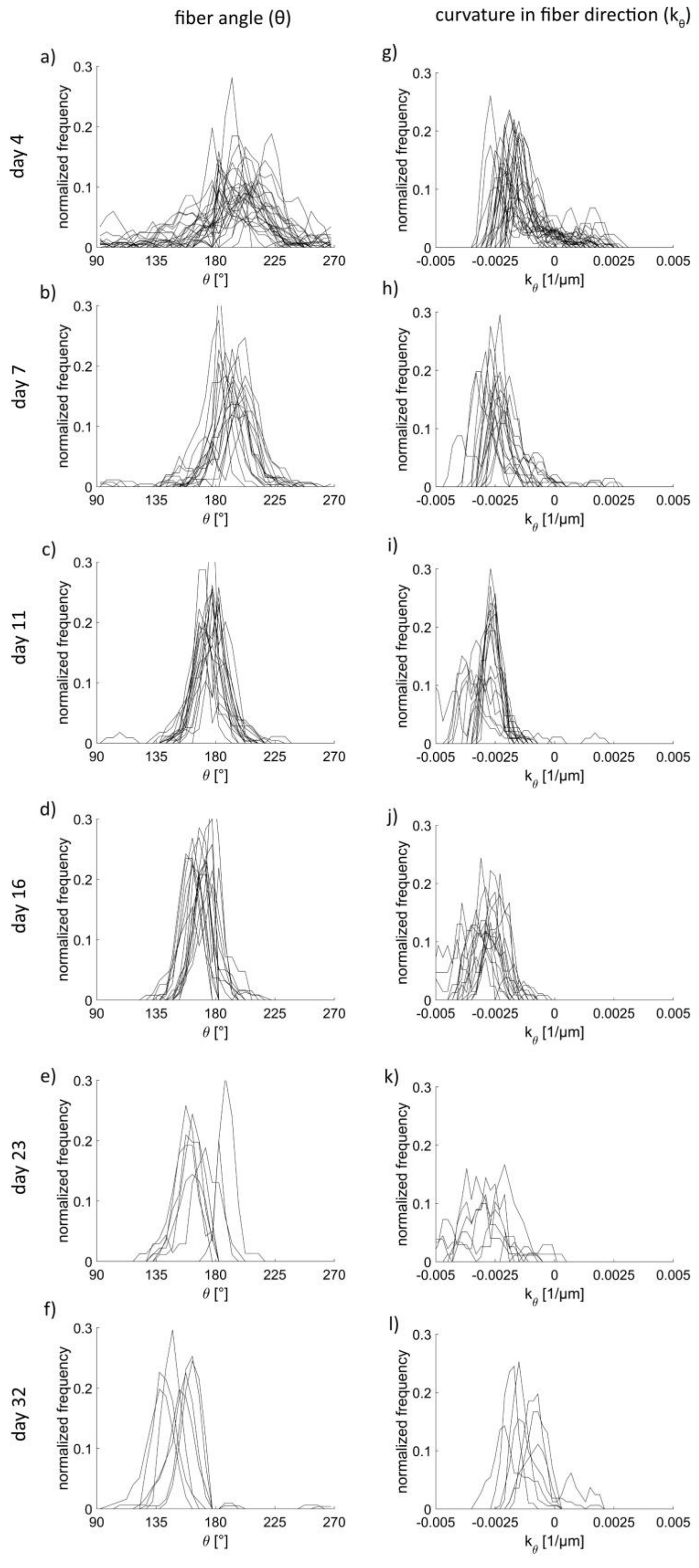


S2: add Fig2: Angle distribution (a-f) for actin and distribution for curvature along theta (g-l) on pores at different timepoints. Each line indicates one sample. $n_{\text{Pore day 4}} = 29$, $n_{\text{Pore day 7}} = 17$, $n_{\text{Pore day 11}} = 17$, $n_{\text{Pore day 16}} = 16$, $n_{\text{Pore day 23}} = 7$, $n_{\text{Pore day 32}} = 7$

### capillary bridges

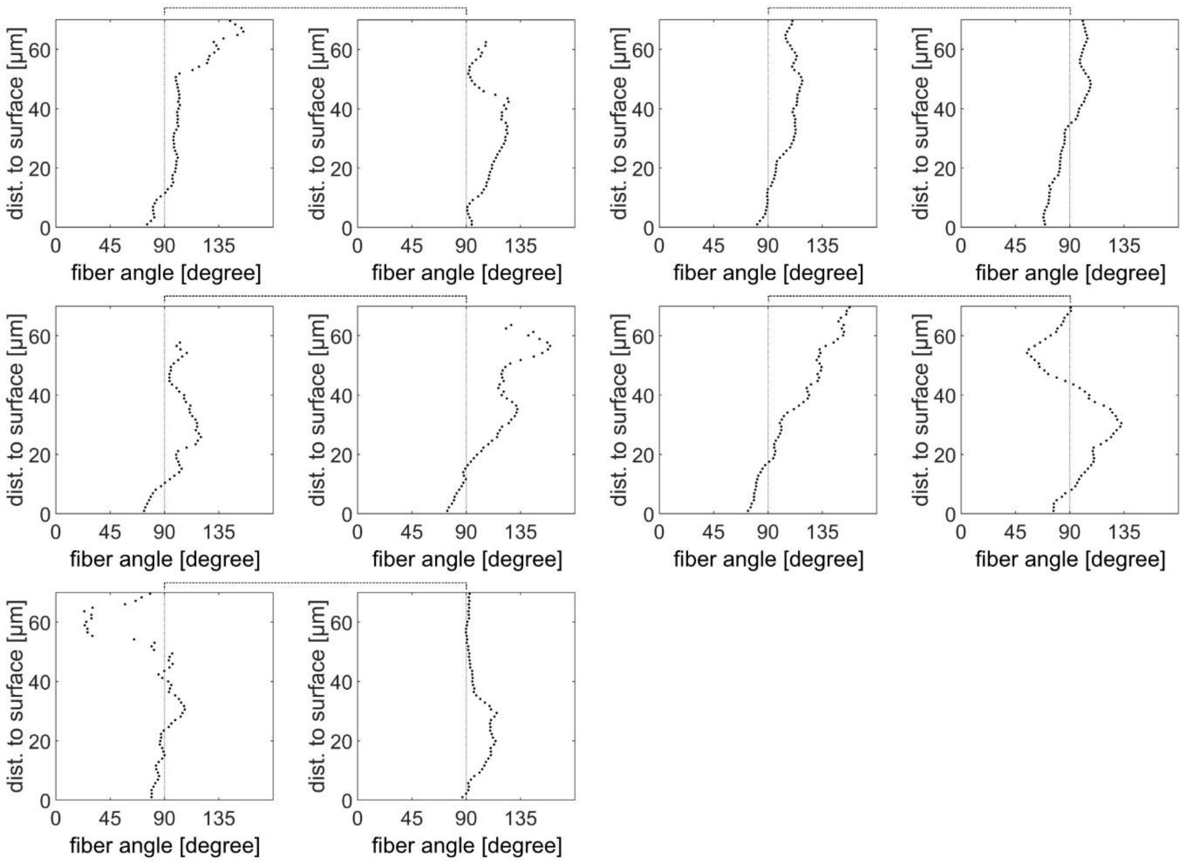


S3: add Fig 3: Evaluation of actin fiber orientations at different depths below the tissue surface in a capillary bridge after 16 days of culture. Each black dot represents the result of a fiber-orientation analysis of a single slice within a region of 200 x 200 µm$^2$. Each plot represents one data set. The connected plots indicate that in these cases two views (180° rotated along the cylinder axis) of one sample were measured.

### pores

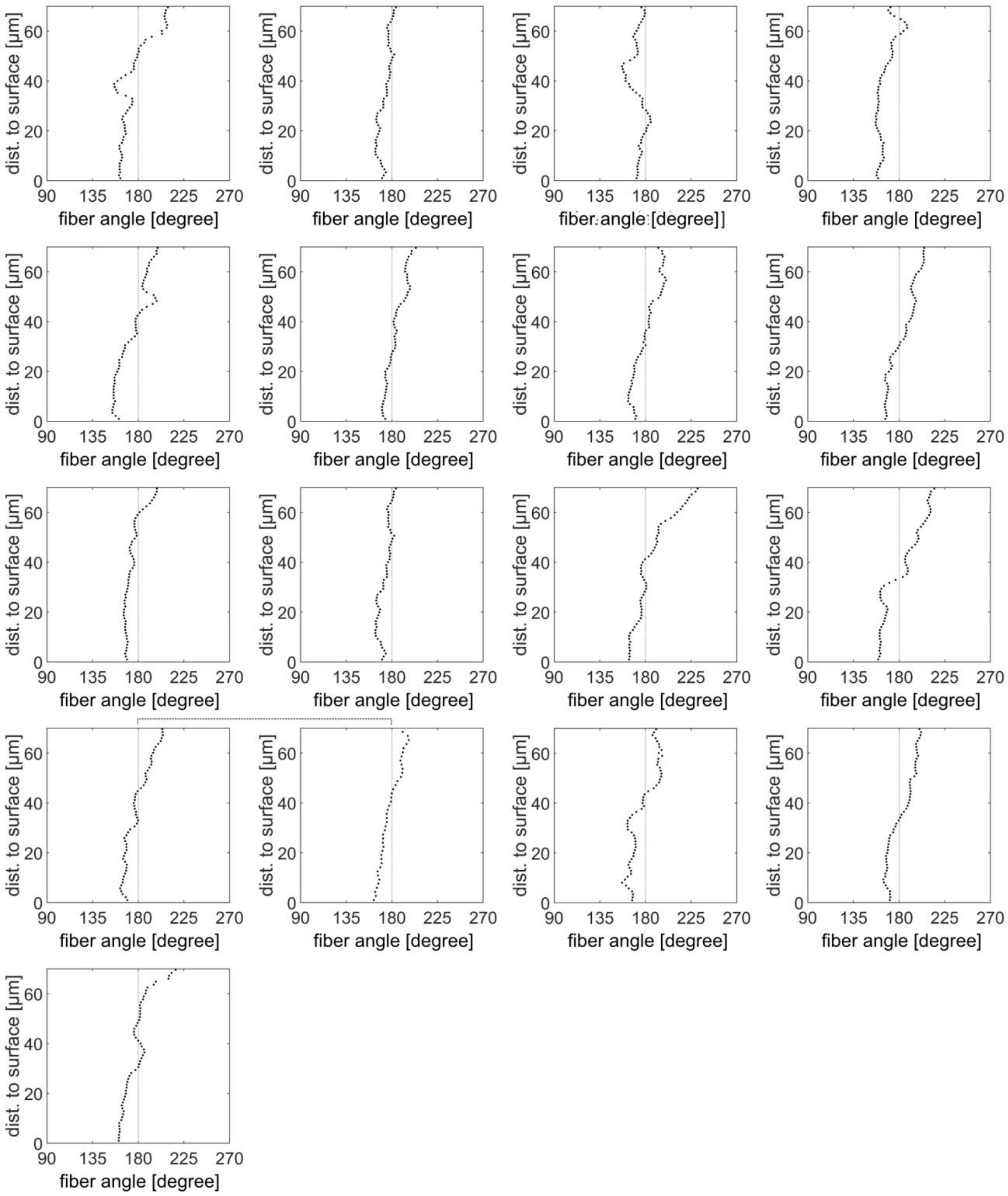


S4: add Fig 3: Evaluation of actin fiber orientations at different depths below the tissue surface in a pore after 16 days of culture. Each black dot represents the result of a fiber-orientation analysis of a single slice within a region of 200 x 200 μm$^2$. Each plot represents one data set. The connected plots indicate that in these cases two views (180° rotated along the cylinder axis) of one sample were measured.

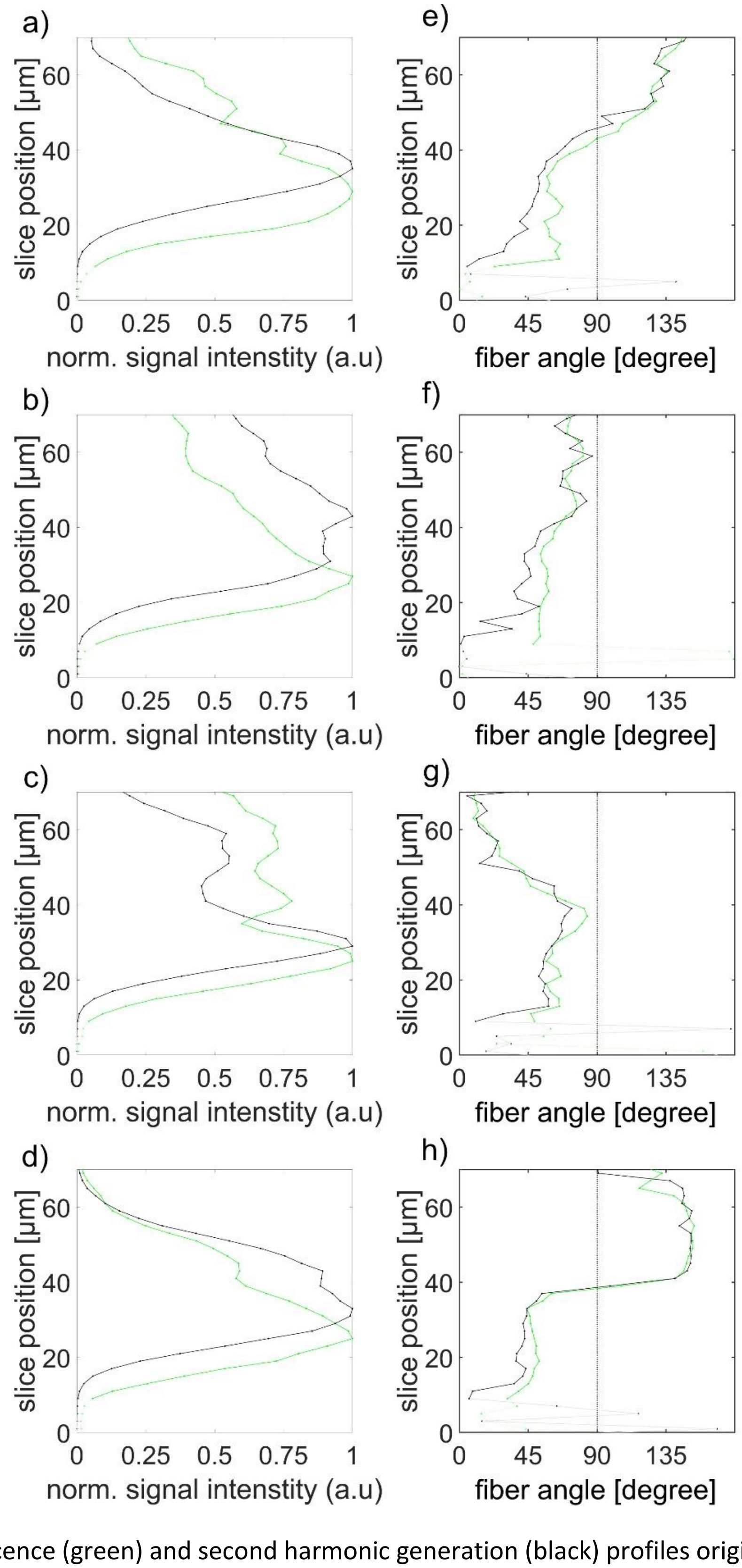


S5: CLSM Fluorescence (green) and second harmonic generation (black) profiles originating from image stacks of capillary bridges. a-c) show the intensity profiles of four samples across the image stack (from the outside 0 µm until 70 µm into the sample); e-h) show the results of the actin and collagen fiber angle evaluations of the same samples. Very low intensity regions are shown in light colours.

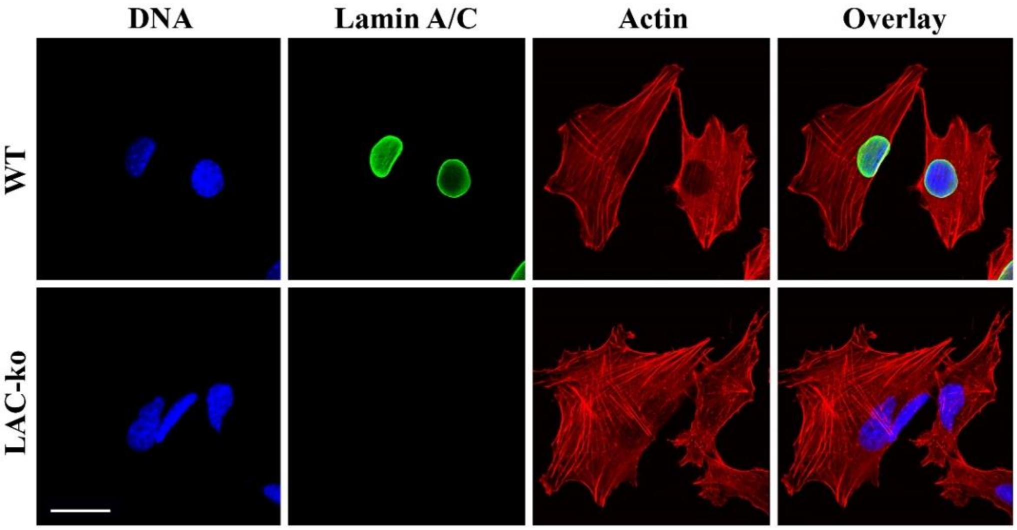


S6: LAC-ko cells lack nuclear lamin A/C. WT and LAC-ko MC3T3-E1 cells were fixed with 4% paraformaldehyde and processed for immunofluorescence microscopy using antibodies against lamin A/C (green). Actin filaments were stained with phalloidin-TRITC (red) and DNA with Hoechst dye (blue). Confocal images are shown (scale bar 20 µm).

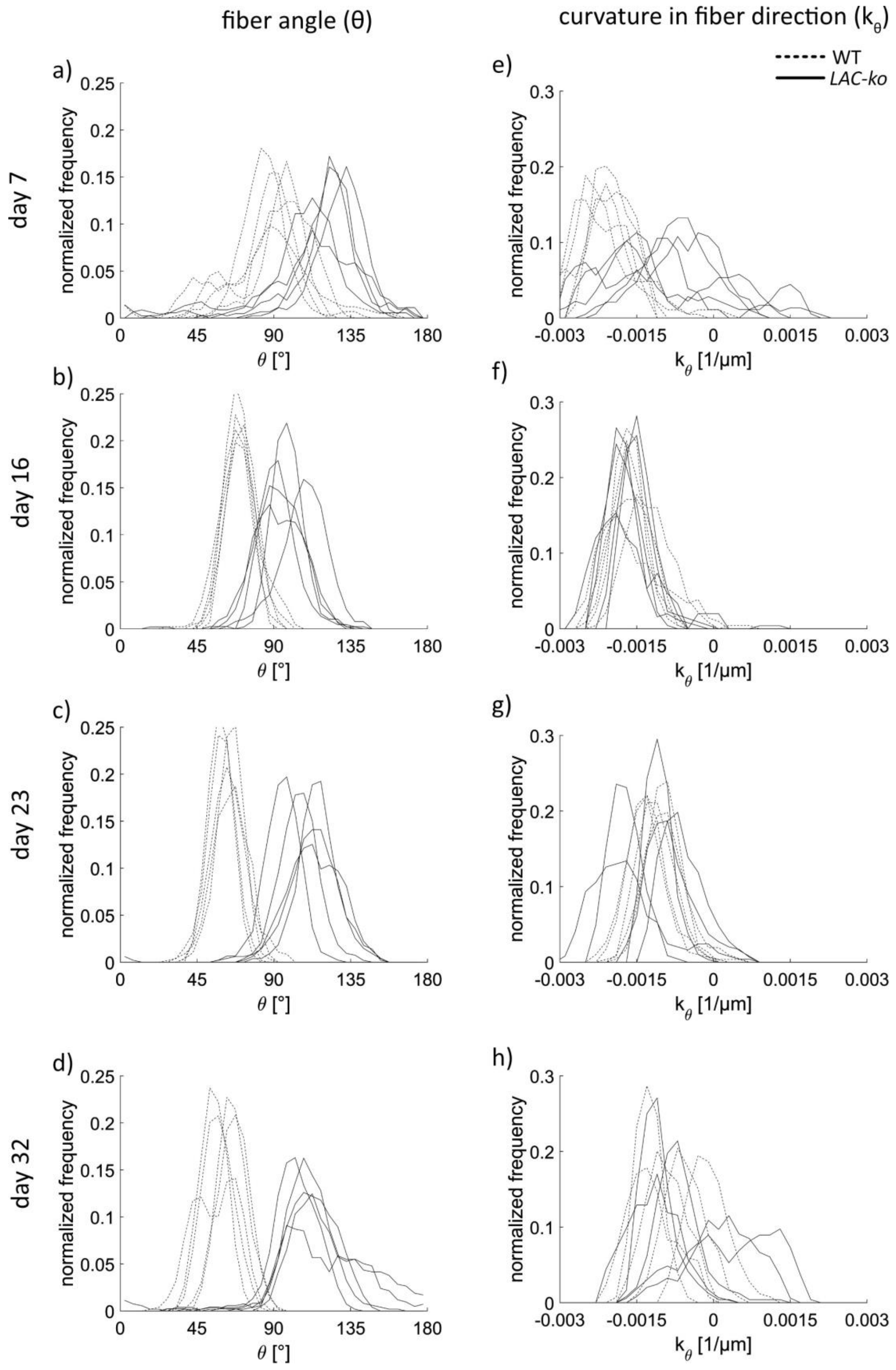


S7: add S7: Fiber alignment of MC3T3-E1 LAC-ko compared to wildtype: Angle distribution (a-d) for actin and distribution for curvature along theta (e-h) on capillary bridges at different timepoints. Each line indicates one sample. Each timepoint consists of five repetitions for WT and LAC-ko.

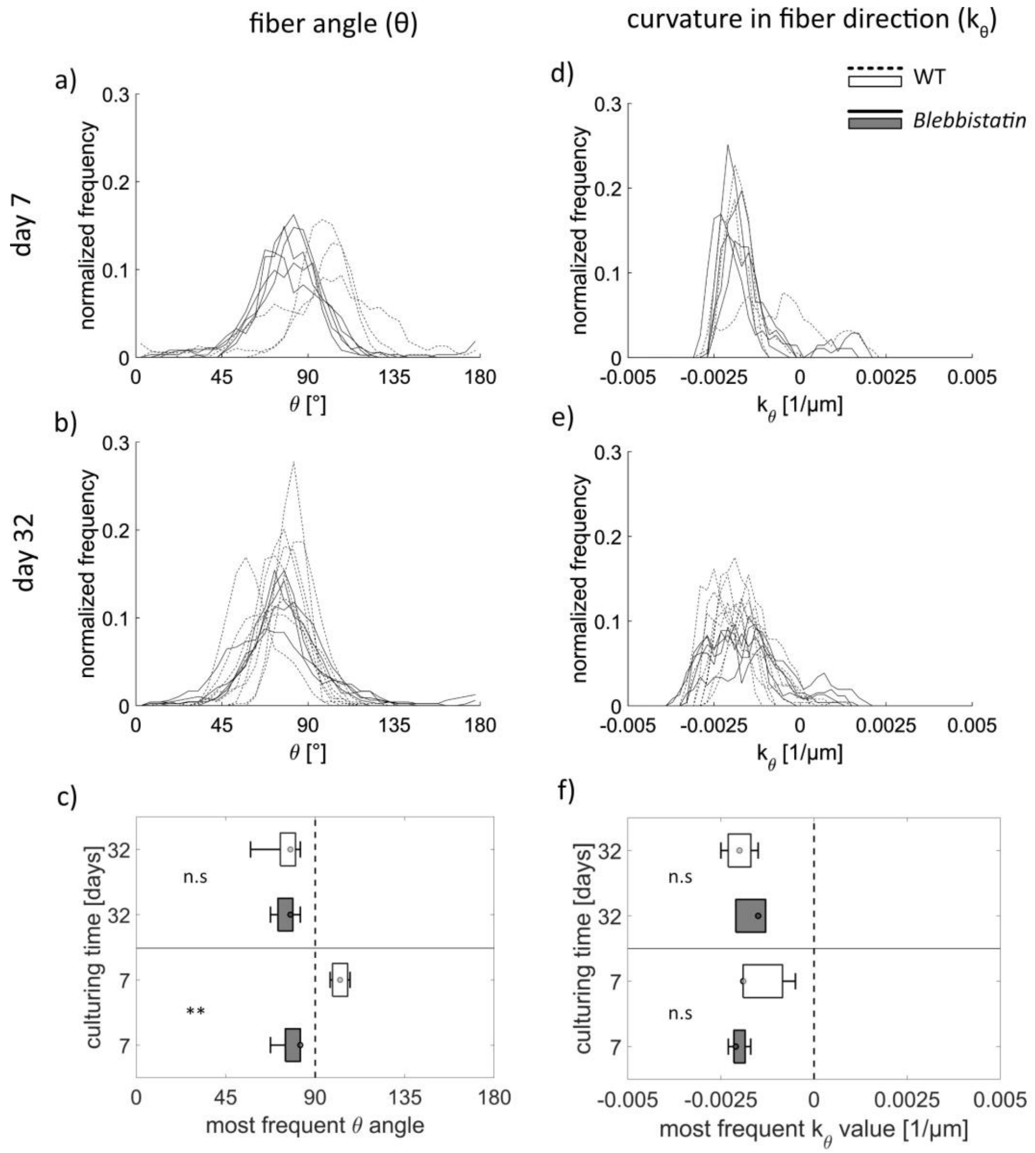


S8: Fiber alignment of MC3T3-E1 compared to samples treated with Blebbistatin: Angle distribution (a, b) for actin and distribution for curvature along theta (d-e) on capillary bridges at day 7 (a, d) or day 32 (b, e). Each line indicates one sample. Average and the standard distribution of the peak position of theta (c) and the curvature along theta (f). $n_{\text{Blebbistatin day 7}} = 5$, $n_{\text{Blebbistatin day 32}} = 5$, $n_{\text{WT day 7}} = 3$, $n_{\text{WT day 32}} = 8$

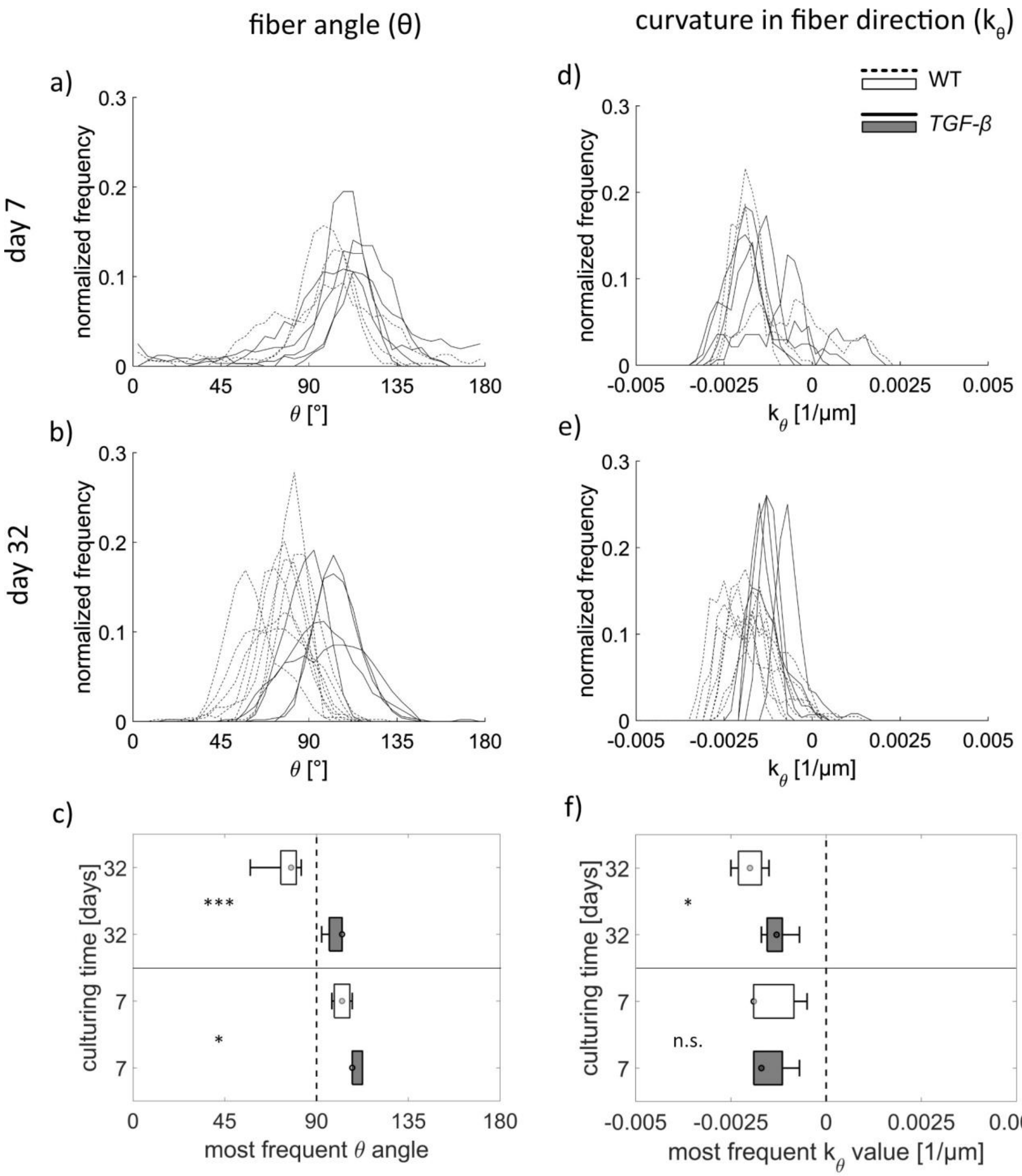


S9: Fiber alignment of MC3T3-E1 compared to samples treated with TGF-b1: Angle distribution (a, b) for actin and distribution for curvature along theta (d-e) on capillary bridges at day 7 (a, d) or day 32 (b, e). Each line indicates one sample. Average and the standard distribution of the peak position of theta (c) and the curvature along theta (f). $n_{TGF\text{-}b1\ day\ 7} = 5$, $n_{TGF\text{-}b1\ day\ 32} = 5$, $n_{WT\ day\ 7} = 3$, $n_{WT\ day\ 32} = 8$

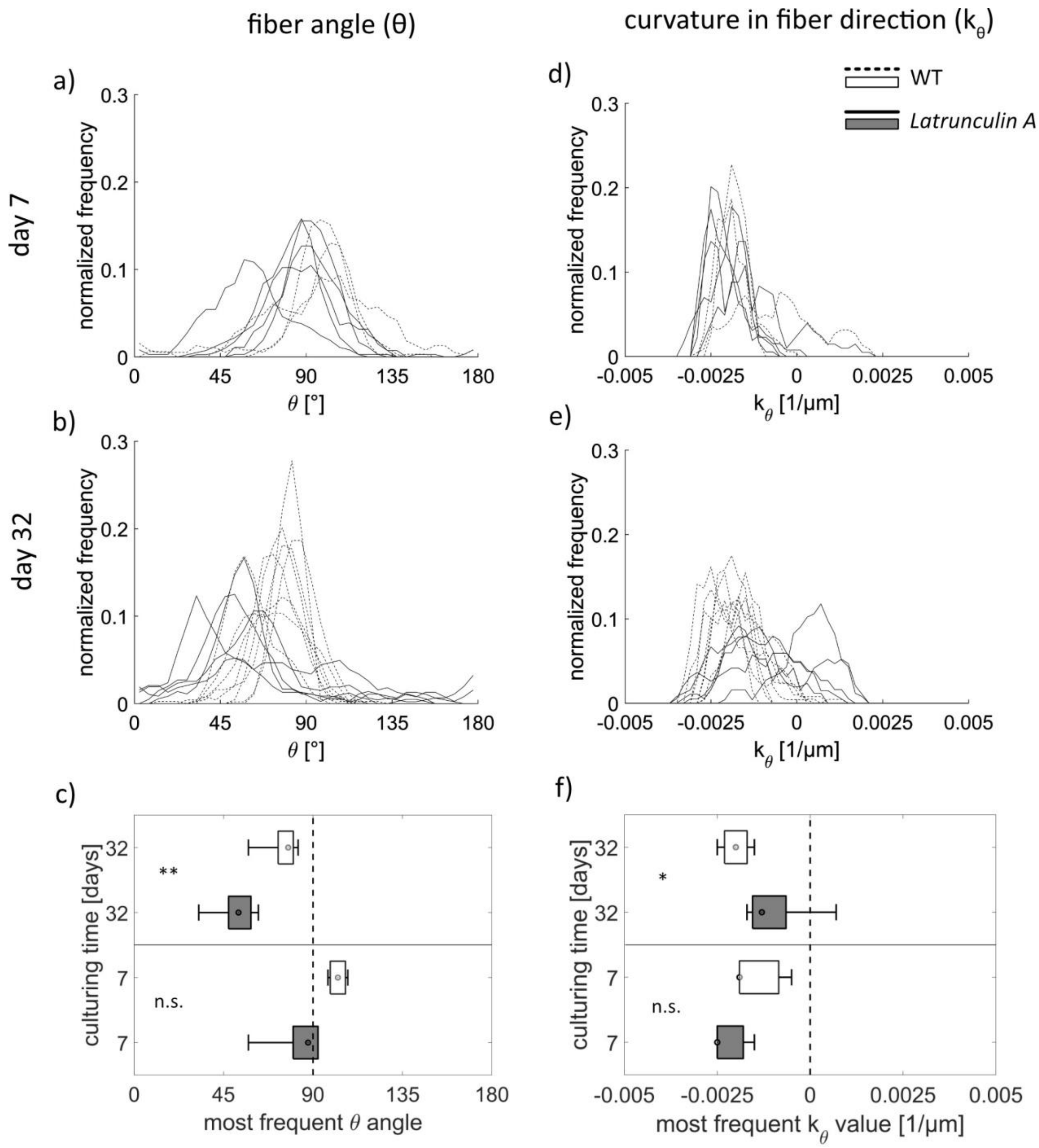


S10: Fiber alignment of MC3T3-E1 compared to samples treated with LatA: Angle distribution (a, b) for actin and distribution for curvature along theta (d-e) on capillary bridges at day 7 (a, d) or day 32 (b, e). Each line indicates one sample. Average and the standard distribution of the peak position of theta (c) and the curvature along theta (f). $n_{LatA\ day\ 7} = 5$, $n_{LatA\ day\ 32} = 5$, $n_{WT\ day\ 7} = 3$, $n_{WT\ day\ 32} = 8$

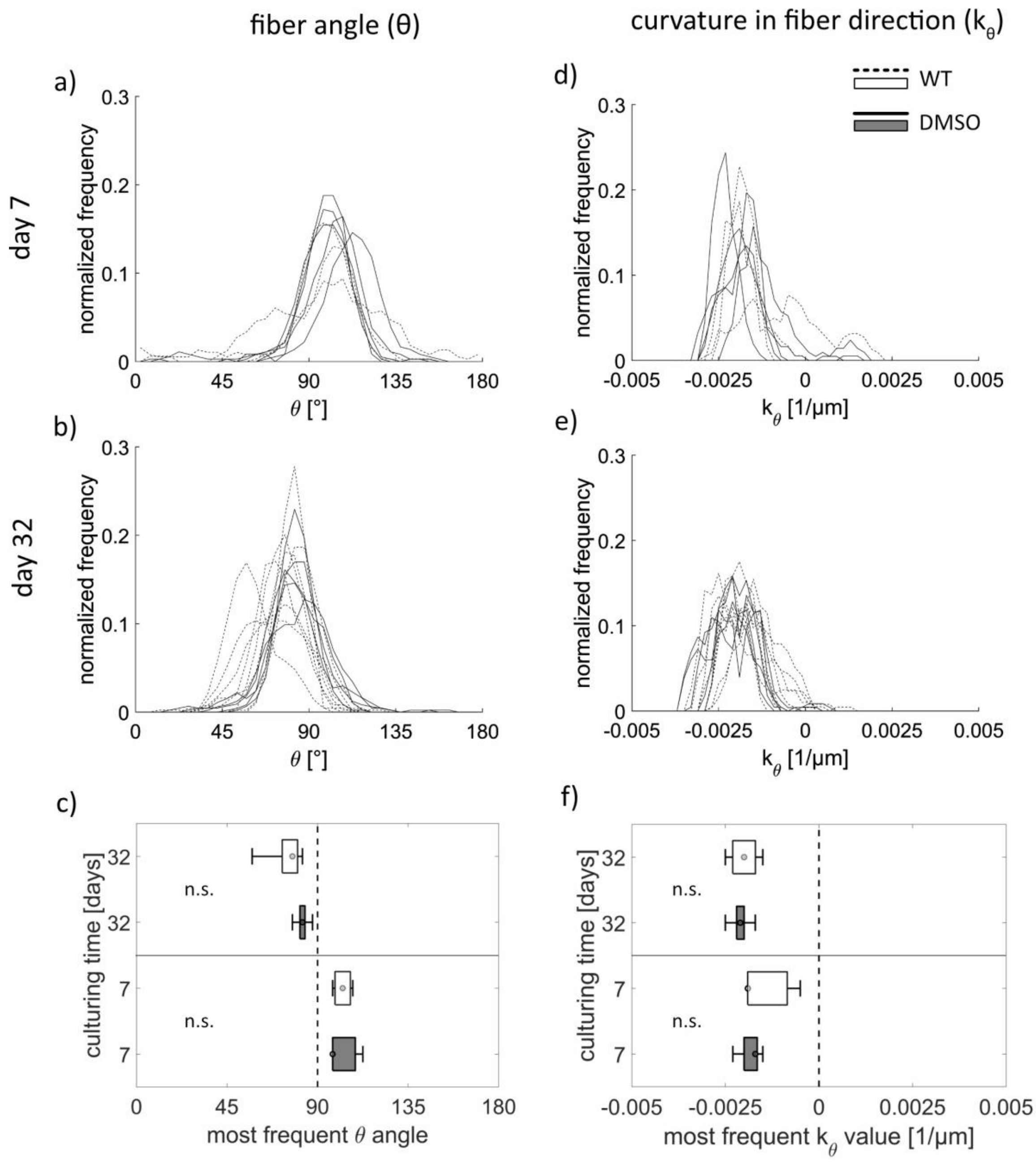


S11: DMSO, Fiber alignment of MC3T3-E1 compared to samples treated with DMSO: Angle distribution (a, b) for actin and distribution for curvature along theta (d-e) on capillary bridges at day 7 (a, d) or day 32 (b, e). Each line indicates one sample. Average and the standard distribution of the peak position of theta (c) and the curvature along theta (f). $n_{DMSO\ day\ 7} = 5$, $n_{DMSO\ day\ 32} = 5$, $n_{WT\ day\ 7} = 3$, $n_{WT\ day\ 32} = 8$

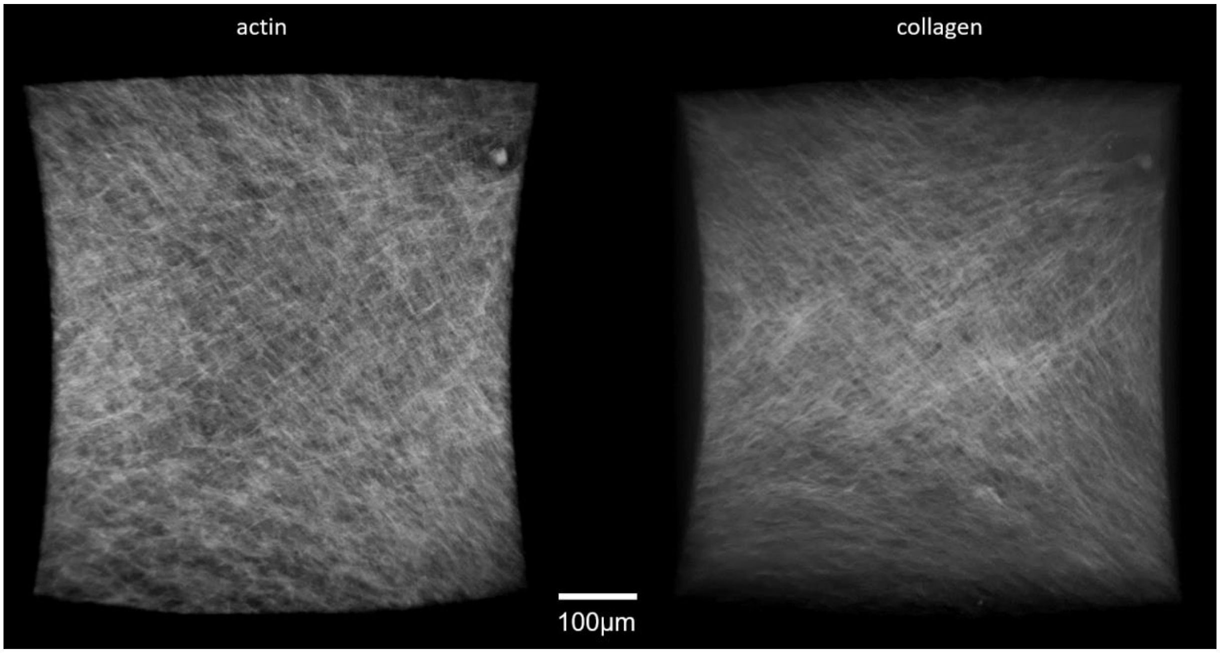


S12: Screenshots from two 360° rotation videos (see supplementary files) of actin and collagen alignment on a tissue surface patch of a capillary bridge sample after 23 days of culturing.

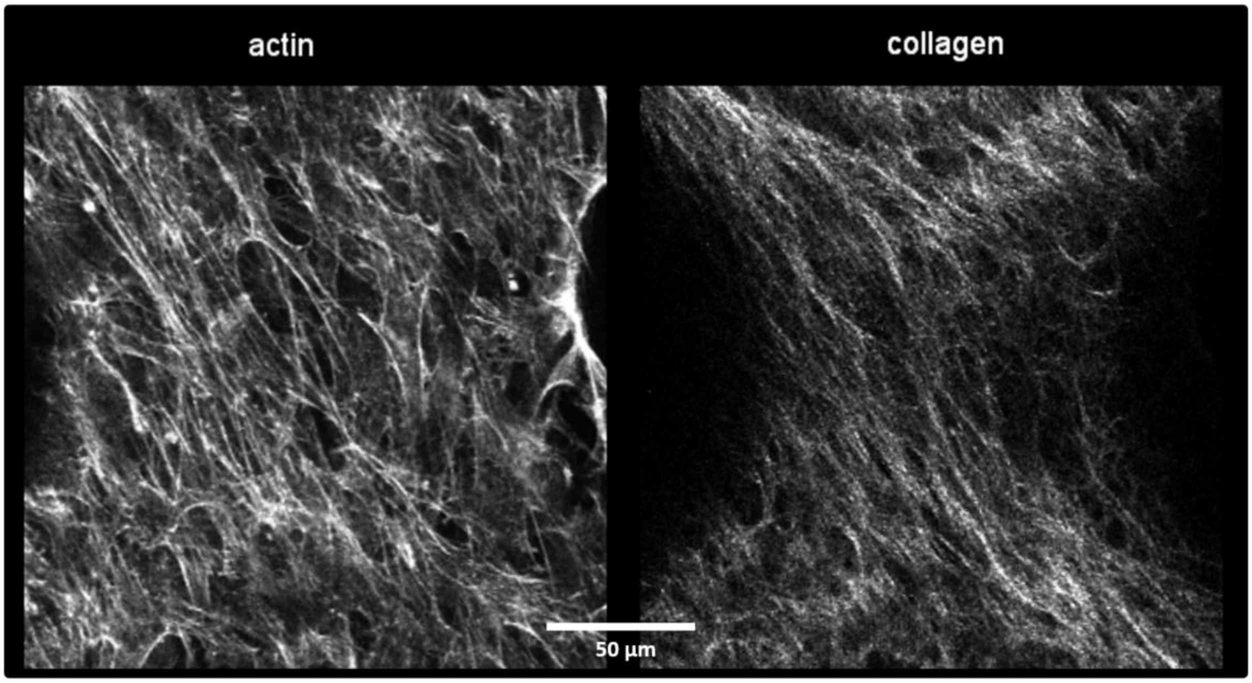


S13: Screenshot from video (see supplementary files) showing depth scan of an actin (CLSM – fluorescence) and collagen (second harmonic generation) of a tissue grown for 16 days on a capillary bridges sample. The video demonstrates the different surface positions, co-alignment of actin and collagen and fiber orientation changes with the distance to the sample surface.

### Materials and Methods

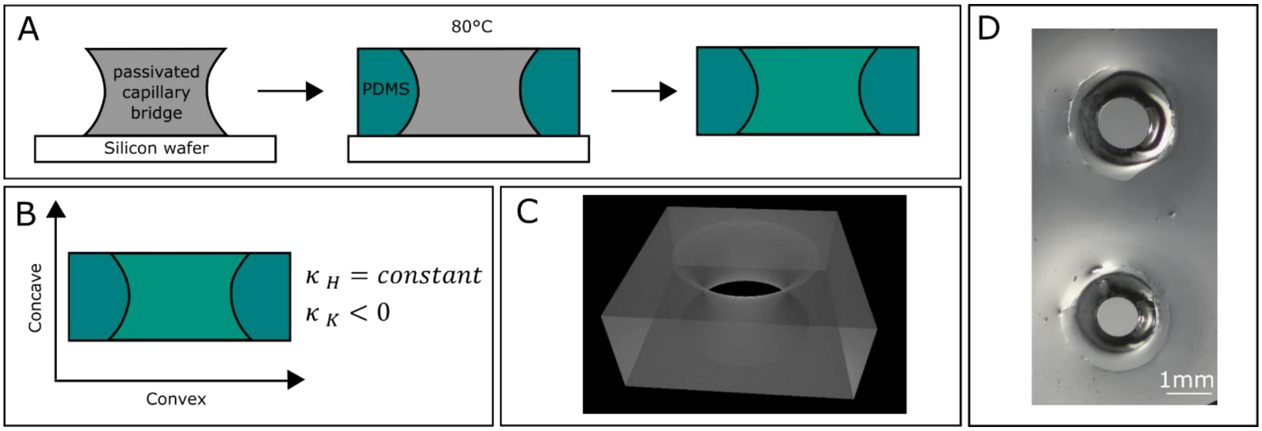


S14: Manufacturing of pores: (A) displays the manufacturing process of a PDMS pore with negative Gaussian curvature surface. (B) Geometrical description of a pore cross-section. (C) Schematic representation of a pore. (D) Representative examples of pores made of PDMS.

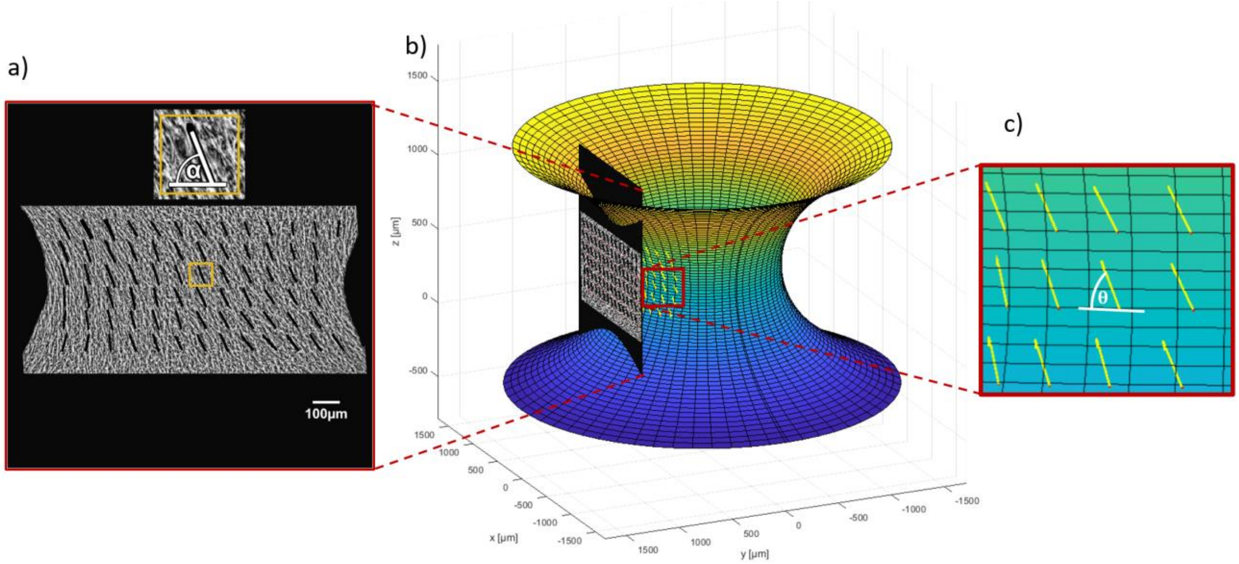


S15: Illustration of the data analysis procedure

a) Maximum intensity projection image of the actin fibers. The projection originates from a z-stack obtained by fluorescence light sheet microscopy after fixing and staining of the sample. Black bars indicate the most prominent local fiber orientation (α) of the respective sub region (81 x 81 µm, orange box) in the projection plane.

b) Vectors representing the fiber direction are projected onto the 3D representation of the sample. The shape of the model is created according to polynomial fits of the sample contours obtained by phase contrast microscopy after fixating the sample.

c) The fiber angle on the curved sample surface relative to the horizontal direction (θ – angle) is derived

**Generation of LAC-ko cells**

MC3T3-E1 cells (Varga, F., Rumpler, M., Klaushofer, K. 1994, FEBS Lett 345(1):67-70) were routinely maintained in α-minimum essential media (α-MEM) (Sigma-Aldrich Chemie GmbH, Steinheim, Germany) supplemented with 4500 mg/L D-(+)-glucose (Sigma-Aldrich Chemie GmbH, Steinheim, Germany), 10% fetal bovine serum (FBS) (gibco, Life Technologies Limited, Paisley, UK), 50 µg/mL L-ascorbic acid (Sigma-Aldrich Chemie GmbH, Steinheim, Germany) and 1% penicillin-streptomycin (Sigma-Aldrich Chemie GmbH, Steinheim, Germany) at 37°C at 5% $CO_2$ in a humidified atmosphere. For CRISPR/Cas9-mediated knockout (1) of lamins A/C, sgRNA sequences were cloned into pSpCas9(BB)-2A-Puro (PX459) V2.0 [gift from Feng Zhang (Addgene plasmid # 62988 ; http://n2t.net/addgene:62988 ; RRID:Addgene_62988)] via BpiI and initially tested in mouse embryonic stem cells at the CRISPR-facility of the Vienna Biocenter Core Facilities (Vienna, Austria) for their ability to edit *LMNA*. One sgRNA (GCAGCCGGGTGATCCGAGT) showing the highest editing efficiency was selected to establish a MC3T3-E1 LAC-ko cell line at the Ludwig Boltzmann Institute of Osteology (Vienna, Austria). In short, MC3T3-E1 cells were transfected with the plasmid coding for the respective sgRNA using the Neon transfection system (Fisher Thermo Scientific, Waltham,MA, USA), cultivated afterwards for 48 h and then seeded into 96-well plates at a density of ~0.5 cells per well. Cell clones growing in individual wells were expanded and analyzed by Sanger sequencing and immunofluorescence microscopy using anti-lamin A/C antibodies (clone 3A6-4C11; Active Motif, Waterloo, Belgium). For sequencing analyses, genomic DNA was isolated and a 1094 bp long region around the site targeted by the sgRNA was amplified by PCR (forward primer: ACCCCCTCCCTTCTATGTCC, reverse primer: GGAAGTGGGGTGAGTCACTG) and sent for sequence analyses (Microsynth, Balgach, Switzerland) using an individual primer (CCTGTAGAGGAGGGCCTATTAGA). Based on the observed editing and the absence of lamin A/C staining one clone was picked and used in this study (LAC-ko cells).

**Immunofluorescence microscopy**

Cells grown on glass coverslips were fixed with 4% paraformaldehyde in PBS for 15 min at room temperature, permeabilized with 0.5% Triton X-100 in PBS and blocked for 30 min in blocking buffer (2% BSA in PBS). Primary anti-lamin A/C antibodies (clone 3A6-4C11; Active Motif, Waterloo, Belgium) were diluted 1:400 in blocking buffer and incubation was performed for 1 h at room temperature. Secondary Alexa Fluor 488 goat anti-mouse IgG was diluted 1:400 in PBS together with phalloidin-TRITC (1:100; Sigma-Aldrich Chemie GmbH, Steinheim, Germany) and Hoechst 33258 (1:10,000; Sigma-Aldrich Chemie GmbH, Steinheim, Germany) and incubated for 30 min at room temperature. Finally cells were mounted in Fluoroshield (Sigma-Aldrich Chemie GmbH, Steinheim, Germany) and images were acquired on a Leica SP5 confocal laser scanning microscope and processed using ImageJ (2).

**Analysis of fiber alignment**

The actin fiber alignment was derived from light sheet microscopy fluorescence data of fixed and stained samples. Due to the anisotropic resolution, instead of using full 3D data, our results are based on maximum intensity projections (Figure S15 a). With respect to the curved 3D sample, fiber angles are distorted in the projection images at all locations except for the neck center region. In order to consider this, first the fiber angles in the MIP image (α) were derived and then projected onto a rotational symmetric 3D representation of the sample, which was created for every sample based on their contour shape (Figure S15 b). In order to obtain the sample contours, bright field images of the capillary bridges before fixation were taken using a Zeiss Axio Vert.A1 microscope with a 5x A-Plan objective. Using in-house developed

Matlab routines (Mathworks Inc., MA, USA, v 2020a), 4[th] order polynomial fits were performed and rotational symmetric representations of the sample were created. Due to the different geometry this was not possible for pore samples, where the contour shape was extracted from the light sheet data. In MIP images obtained from the light sheet microscope, obvious artefacts were masked to be excluded from the further analysis before the image was sub-divided into boxes of 81 x 81 µm using in-house developed Matlab scripts. For each of this box a fast fourier transform was performed followed by filtering and thresholding and the fit of an ellipse to obtain the main fiber orientation and thus the α angle. The routines for the fiber angle alignment are based on the work published in reference (3). Edge regions as well as regions more than 300 µm above/below the neck center were excluded from the evaluation to avoid boundary artefacts. Subsequently, α angles were projected onto the 3D representation of the sample thus obtaining the $\theta$-angle which now represents the actual actin fiber angle on the surface as shown in Figure 1 of the main manuscript and SI Figure 15. For each light sheet measurement the $\theta$-angle was derived on around 100 different locations being the basis for the presented histograms.
